# Expanding magnetic organelle biogenesis in the domain *Bacteria*

**DOI:** 10.1101/2020.04.27.061960

**Authors:** Wei Lin, Wensi Zhang, Greig A. Paterson, Qiyun Zhu, Xiang Zhao, Rob Knight, Dennis A. Bazylinski, Andrew P. Roberts, Yongxin Pan

## Abstract

The discovery of membrane-enclosed, metabolically functional organelles in *Bacteria* and *Archaea* has transformed our understanding of the subcellular complexity of prokaryotic cells. However, whether prokaryotic organelles emerged early or late in evolutionary history remains unclear and limits understanding of the nature and cellular complexity of early life. Biomineralization of magnetic nanoparticles within magnetosomes by magnetotactic bacteria (MTB) is a fascinating example of prokaryotic organelles. Here, we reconstruct 168 metagenome-assembled MTB genomes from various aquatic environments and waterlogged soils. These genomes represent nearly a 3-fold increase over the number currently available, and more than double the known MTB species. Phylogenomic analysis reveals that these newly described genomes belong to 13 Bacterial phyla, six of which were previously not known to include MTB. These findings indicate a much wider taxonomic distribution of magnetosome organelle biogenesis across the domain *Bacteria* than previously thought. Comparative genome analysis reveals a vast diversity of magnetosome gene clusters involved in magnetosomal biogenesis in terms of gene content and synteny residing in distinct taxonomic lineages. These gene clusters therefore represent a promising, diverse genetic resource for biosynthesizing novel magnetic nanoparticles. Finally, our phylogenetic analyses of the core magnetosome proteins in this largest available and taxonomically diverse dataset support an unexpectedly early evolutionary origin of magnetosome biomineralization, likely ancestral to the origin of the domain *Bacteria*. These findings emphasize the potential biological significance of prokaryotic organelles on the early Earth and have important implications for our understanding of the evolutionary history of cellular complexity.

## Introduction

It was accepted widely that intracellular, membrane-bounded, metabolically functional organelles are present exclusively in eukaryotic cells and that they are absent from *Bacteria* and *Archaea*. This long-held view was revised after numerous recent discoveries of a diverse group of highly organized, membrane-enclosed organelles in the domains *Bacteria* and *Archaea* associated with specific cellular functions ^1–3^. However, the origin and evolution of prokaryotic organelles remain largely elusive. It is still unclear whether organelle biogenesis emerged early or late during the evolution of *Bacteria* and *Archaea*, posing problems for elucidating the evolutionary history of cellular complexity.

Magnetosomes within magnetotactic bacteria (MTB) are a striking example of prokaryotic organelles ^4^. Magnetosomes consist of a lipid bilayer-bounded membrane in which nanosized, ferrimagnetic magnetite (Fe_3_O_4_) and/or greigite (Fe_3_S_4_) crystals are biomineralized and usually arranged in chain-like structure(s) that maximize the magnetic dipole moment ^5,6^ (Figure 1). The most accepted major function of magnetosomes is to produce tiny compass needles that facilitate MTB navigation to their preferred low-O_2_ or anaerobic microenvironments in chemically-stratified aquatic systems, a behaviour referred to as magnetotaxis or microbial magnetoreception ^7^. Additional suggested functions of magnetosomal crystals include detoxification/elimination of toxic reactive oxygen species (ROS), iron sequestration and storage in which they act as an electrochemical battery, or as a gravity sensor ^8–10^ (Figure 1). Understanding the phylogenetic and genomic diversity of MTB could advance significantly our understanding of the evolutionary origin of bacterial organelle biogenesis in general. Moreover, considering that magnetoreception occurs widely in both micro- and macro-organisms and that magnetosomal crystals are the only magnetoreceptors definitively characterized thus far, MTB also represent a valuable system for exploring the origin and early evolution of magnetoreception ^11–14^.

**Figure 1.**
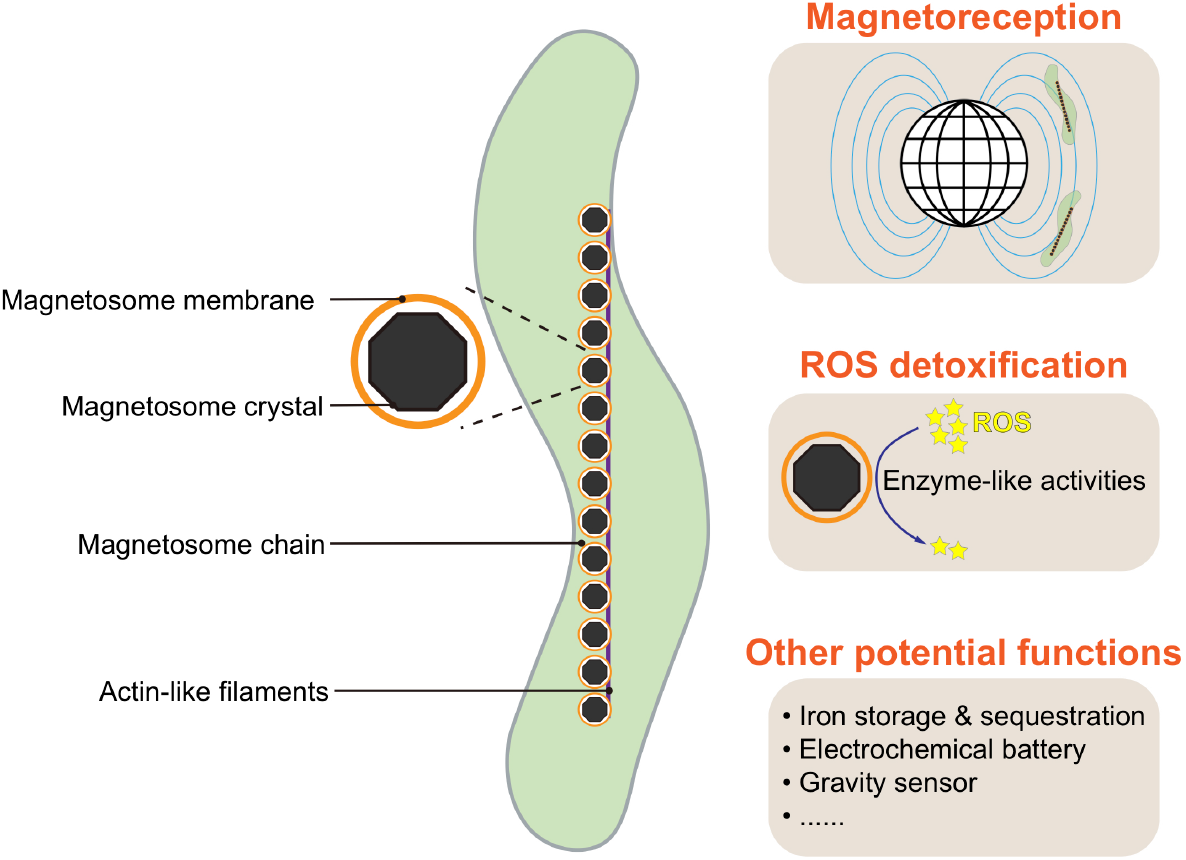
The magnetosome in which magnetotactic bacteria (MTB) biomineralize magnetic crystals is a typical example of a bacterial organelle. Membrane-bounded magnetosomes contain intracellular magnetic nanoparticles (Fe_3_O_4_ or Fe_3_S_4_), with typical ~20-150 nm sizes. Magnetic particles within MTB magnetosomes are typically organized into (a) chain-like structure(s) within the cell in order to optimize the cellular magnetic dipole moment. Functions of magnetosomes include magnetoreception ^103,104^ and ROS detoxification ^78,79^, both of which have been experimentally proven. Additional proposed functions, such as iron storage and sequestration, acting as an electrochemical battery or a gravity sensor, need further testing.

MTB are distributed globally across a broad range of O_2_-limited or anaerobic aquatic habitats, ranging from freshwater lakes to oceans and even to some extreme environments ^15–17^. However, few studies have reported MTB in waterlogged soils ^18,19^. Magnetosomal biogenesis by MTB is recognized as a key component of the global iron cycle ^20,21^, and is also potentially important in the biogeochemical cycling of phosphorus, nitrogen, and sulphur ^9,22–24^. Historically, magnetosomal biomineralization has been viewed as a specialized type of metabolism restricted to two Bacterial phyla: the *Proteobacteria* (*Alphaproteobacteria*, *Deltaproteobacteria*, and *Gammaproteobacteria* classes) and the *Nitrospirae* ^25^. Initial attempts to explain this restricted yet scattered distribution of magnetosomal biogenesis based on a few taxa gave rise to two alternative hypotheses: polyphyletic origin in different taxonomic lineages ^26^, or extensive horizontal gene transfers (HGTs) ^27,28^.

In recent years, the known extent of MTB diversity has undergone a significant expansion due to methodological advances, such as the successful cultivation of novel MTB strains ^e.g., 29–32^, 16S rRNA gene-based characterization ^e.g., 33–35^, genome mining of public repositories including GenBank and IMG/ER ^17,36^, and cultivation-independent surveys of magnetosome gene clusters (MGCs, which are physically clustered groups of genes that are together responsible for magnetosomal biogenesis) ^37,38^. MTB have now been further identified in other Bacterial taxa, including the *Betaproteobacteria*, *Zetaproteobacteria*, “*Candidatus* Etaproteobacteria”, and “*Candidatus* Lambdaproteobacteria” classes of the *Proteobacteria* phylum, the candidate phylum *Omnitrophica* (previously known as the candidate division OP3), the candidate phylum *Latescibacteria* (previously known as the candidate division WS3), and the phylum *Planctomycetes*, according to the NCBI taxonomy. Analyses including data from these latter groups suggest that magnetosomal biogenesis among different Bacterial lineages has a monophyletic origin from a common ancestor, which occurred prior to divergence of the *Nitrospirae* and *Proteobacteria* phyla, or perhaps even earlier, in the last common ancestor of five MTB-containing Bacterial phyla: *Proteobacteria*, *Nitrospirae*, *Omnitrophica*, *Latescibacteria*, and *Planctomycetes* ^15,37,39–41^.

Discovery of novel MTB outside the traditionally recognized taxonomic lineages suggests that the diversity and distribution of MTB across the domain *Bacteria* are considerably underestimated. This raises important questions regarding the origin and evolution of magnetosome organelle biosynthesis. Here we present the most comprehensive metagenomic analysis available of MTB communities from geographically, physically, and chemically diverse sites to evaluate two questions: (1) are MTB taxonomically widespread among the phyla of the domain *Bacteria*; and (2) did the magnetosome organelle originated early?

## Results and Discussion

### Survey of MTB from diverse environments

We perform a light microscopy survey of MTB from wide-ranging environments across China and Australia (Figure 2a), including sediments from freshwater lakes, ponds, rivers, creeks, paddy fields, intertidal zones, and soils from acidic peatlands. MTB were observed microscopically in these habitats with a salinity range of <0.1-37.0 ppt and a pH range of 4.3-8.6 (Supplementary Table 1). Unexpectedly, we find living MTB cells in acidic peatland soils (pH 4.3-5.7) with high water contents (>60%) and organic matter contents (typically >20%) ^42–44^. MTB have been found broadly in diverse aquatic ecosystems ^17^, including some extreme environments such as hot springs ^45,46^, saline-alkaline lakes ^47^, acidic lagoons and mine drainage systems ^33,48^, and deep-sea sediments ^49^, but reports of MTB in soils are limited to a few studies that were published almost 30 years ago ^18,19^. Whether MTB can survive and reproduce in waterlogged soil environments remains unresolved, and the taxonomic diversity of MTB in these environments has not been elucidated. Our finding represents, to the best of our knowledge, the first discovery of MTB in soils with relatively acidic pH. MTB in acidic peatland soils are represented mainly by deep-branching lineages, which is markedly different from other environments (Figure 3, discussed below). Our finding, together with the previous studies ^18,19^, suggests widespread MTB occurrence in different types of waterlogged soils, in addition to subaqueous sediments and bodies of water.

**Figure 2.**
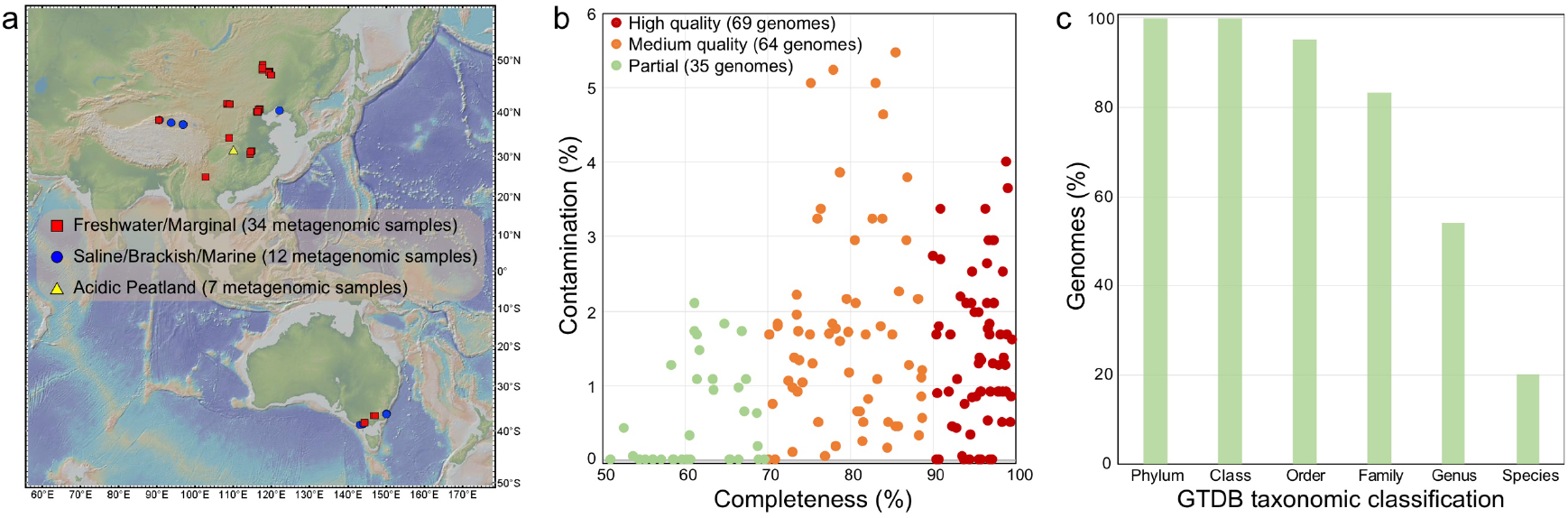
Recovery of 168 MTB genomes from various environments. **a.** Map of sampling locations (generated using the GeoMapApp 3.6.0, http://www.geomapapp.org/). Further site details are given in Supplementary Table 1. **b.** Estimated completeness and contamination of MTB genomes reconstructed in this study. CheckM was used to estimate completeness and contamination. Of these genomes, 69 are high-quality (>90% completeness and <5% contamination), 64 are medium-quality (>70% completeness and <10% contamination) and 35 are partial (50-70% completeness and <5% contamination) genomes. **c.** Relative abundance of recovered MTB genomes that can be classified according to the GTDB taxonomy (database Release 04-RS89). Of the 168 recovered genomes, 34 were classified at species level, 91 were classified at genus level, 140 were classified at family level, 160 were classified at order lever, and 168 could be classified at class and phylum levels. Details are given in Supplementary Table 2.

**Figure 3.**
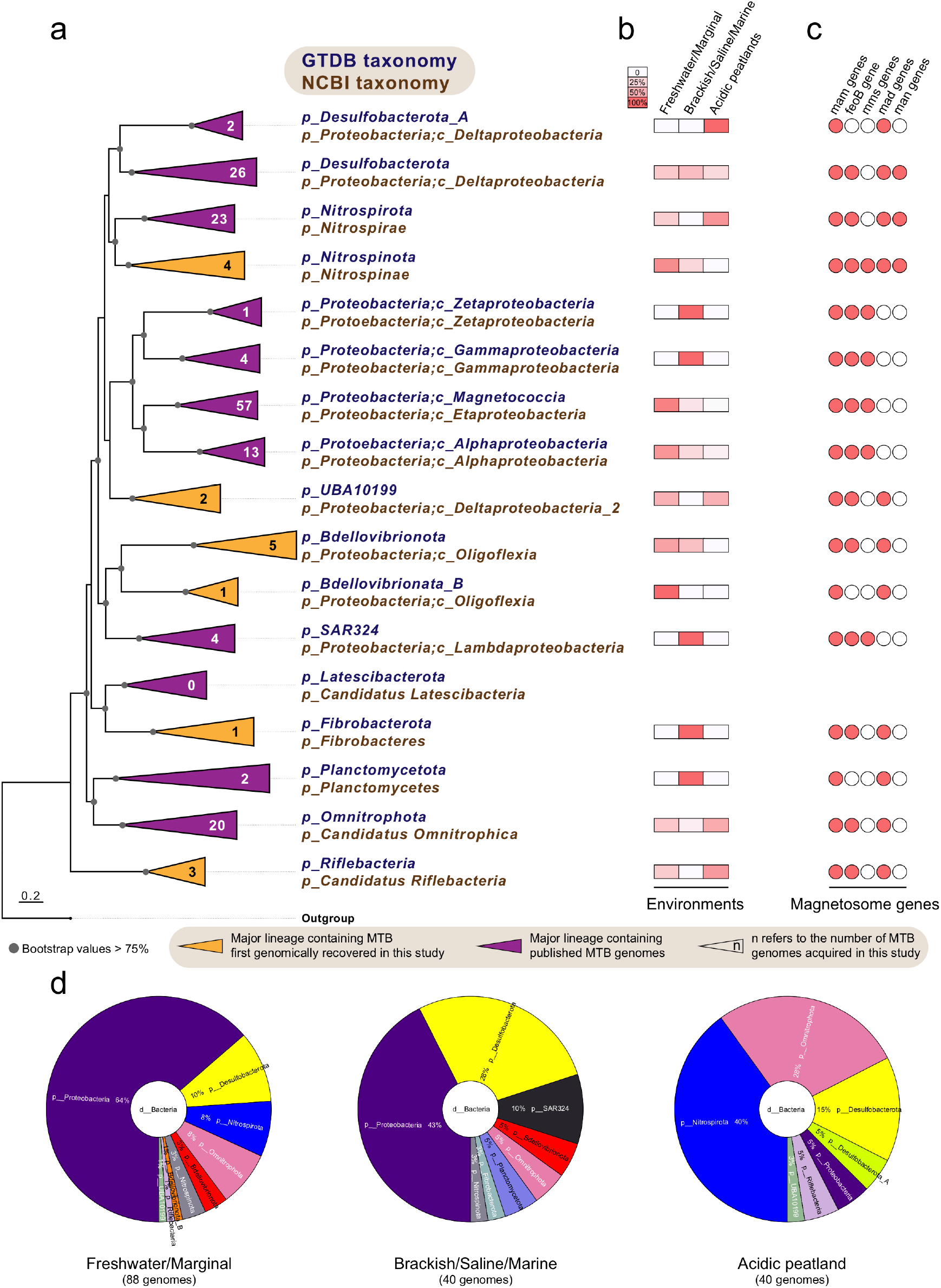
Distribution of MTB genomes across Bacterial phyla and distinct environments. **a.** The maximum-likelihood phylogenomic tree of MTB genomes and their close non-MTB relatives inferred from concatenated 120 Bacterial single-copy marker proteins ^50^, which was constructed using IQ-TREE under the LG+I+G4 substitution model. The number in each clade refers to the number of MTB genomes reconstructed in this study. The complete tree is shown in Supplementary Figure 1. **b.** Relative abundances of reconstructed MTB genomes in this study across different environments within each lineage. **c.** Distribution of magnetosome genes (*mam*, *mms*, *mad*, and *man*) and *feoB* gene within MGCs across different lineages. **d.** Distribution of acquired MTB genomes at the phylum level across different environments, including freshwater/marginal (<1 ppt) and saline/brackish/marine (>1 ppt) sediments, and soils from acidic peatland. Details are given in Supplementary Table 1.

### Expanded genomic diversity of MTB

Metagenomic DNA from magnetically enriched MTB was sequenced and metagenome-assembled genomes were reconstructed with a single-sample assembly and binning strategy (see Methods). Reconstructed genomes were checked manually for the presence of MGCs. From this analysis, we recover a total of 168 MGC-containing genomes with a quality score ^50^ above 50 (Supplementary Table 2). Of these putative MTB genomes, 69 (41%) are high-quality (with >90% completeness and <5% contamination), 64 (38%) are medium-quality (>70% complete with <10% contamination), and 35 (21%) are partial (with 50-70% completeness and <5% contamination) genomes (Figure 2b and Supplementary Table 2). These genomes increase substantially the known genomic diversity of MTB, expanded from 59 (as of January 2020, Supplementary Table 3) to 227 MTB genomes, which can now be clustered into 164 unique species-level genomes based on a 95% average nucleotide identity (ANI) ^51^. Of these species, 110 (67%) are reported here for the first time.

The taxonomy of the newly acquired MTB genomes is classified using a standardized phylogenomic curated taxonomy system Genome Taxonomy Database (GTDB) Toolkit ^52,53^. Around 80% of the recovered genomes cannot be assigned at the species level, and more than 45% cannot be assignable at the genus level (Figure 2c), which confirms that many of these genomes represent previously unknown populations. The phylogenetic relationships of these genomes were further determined by phylogenomic analysis (see Methods). The reconstructed genome tree is congruent with the GTDB taxonomy as shown in Figure 3a and Supplementary Figure 1; therefore, the GTDB taxonomy is used for taxonomic classification of MTB throughout, unless otherwise noted. The corresponding NCBI taxonomy of each lineage is also given in Figure 3a.

The 168 genomes belong to organisms from 13 distinct Bacterial phyla as defined in the GTDB taxonomy and 7 phyla according to the NCBI taxonomy (Figure 3a and Supplementary Table 2). These genomes are recovered from several previously poorly characterized MTB groups, including 26 genomes from the *Desulfobacterota* phylum, 23 genomes from the *Nitrospirota* phylum, and 20 genomes from the *Omnitrophota* phylum (Figure 3a). These novel genomes also expand substantially the representation of common MTB lineages, such as the *Magnetococcia* (57 genomes) and *Alphaproteobacteria* (13 genomes) classes of the *Proteobacteria* phylum. More importantly, we identify 16 genomes that are affiliated within 6 phyla that were not known previously to contain MTB, including the *Nitrospinota*, UBA10199, *Bdellovibrionota*, *Bdellovibrionata*_B, *Fibrobacterota*, and *Riflebacteria* phyla (Figure 3a and Supplementary Figure 1). This expands greatly the number of Bacterial lineages associated with magnetosomal biogenesis and magnetic navigation.

To deepen our understanding of the environmental distribution patterns of MTB, the taxonomic diversity of novel MTB genomes was compared across environments. Genomes belonging to the phyla *Proteobacteria* and *Desulfobacterota* are found predominantly in both freshwater/marginal (<1 ppt) and saline/brackish/marine (>1 ppt) sediments, whereas genomes affiliated with the *Nitrospirata* and *Omnitrophota* phyla represent the dominant MTB groups in acidic peatland soils (Figure 3d). MTB from the *Proteobacteria*, *Desulfobacterota*, and *Omnitrophota* exist in all three environmental sample types, while those of the *Desulfobacterota*_A phylum and of the SAR324, *Fibrobacterota*, and *Planctomycetota* phyla are observed exclusively in acidic peatland soils and brackish/saline/marine environments, respectively (Figure 3b). For *Proteobacteria* classes, MTB genomes from the *Magnetococcia* and *Alphaproteobacteria* are found in wide-ranging environments, while those with *Zetaproteobacteria* and *Gammaproteobacteria* genomes occur solely in brackish/saline/marine environments.

### Diverse MGCs across distinct taxonomic lineages

Genes for the metabolic pathway responsible for magnetosomal biogenesis have been found in contiguous gene clusters in MTB genomes ^54^, which are referred to as MGCs. These gene clusters are not only the key to deciphering the mechanisms and evolutionary origin of magnetosome formation and magnetotaxis ^5^, but they also provide a wealth of gene resources for biosynthesis of membrane-bounded, single-domain magnetic nanoparticles with diverse properties for various applications ^55^. The genomes acquired here contain diverse MGC types in terms of gene content and synteny (Figure 4), which expands our knowledge of MGC diversity considerably. Discovery of various MGCs suggests the potential for diverse magnetosomal biogenesis and magnetotaxis across the domain *Bacteria*. The *mam* ^56–58^ genes that play essential roles in magnetosomal biogenesis are present in all MGCs identified here. Remarkably, we note that the *feoB* gene, which is responsible for iron transport into the cell, is also shared by most MTB lineages and is usually included in the MGCs (Figure 3c and Figure 4). Deletion of *feoB* in *Magnetospirillum* strains results in reduced magnetite biomineralization ^59,60^, which indicates its potentially significant role in magnetosomal biomineralization. The *mad* ^61,62^ and *man* ^63^ genes, which have been proposed to play important accessary functions in magnetosomal biomineralization, represent a much wider distribution across MTB genomes than previously thought: *mad* genes are present in the genomes of 11 phyla and *man* genes occur in the genomes of the *Desulfobacterota* (nTS_bin18 and nDJH15_bin4), *Nitrospirota*, and *Nitrospinota* (nPCR_bin9 and nNGH_bin12) phyla. The *mms6* operon, which contains magnetosome genes *mms6*, *mmsF*, *mms36*, and *mms48*, controls the size and/or number of magnetic magnetosomal crystals in *Magnetospirillum* strains ^57,58,64,65^, and these genes have previously only been found in the *Proteobacteria* phylum. Here we find that MGCs from genomes of the phyla *Nitrospinota* (nNGH_bin12) and SAR324 (nKLK_bin6 and nPCR_bin7) also contain *mms* genes (*mms6* and *mmsF*). In general, the gene content and orientation of MGCs vary considerably across different taxonomic lineages but are generally conserved within the same lineage (Figure 4), which is indicative of lineage specific MGC evolution without extensive inter-phylum HGTs and inter-class HGTs among the *Proteobacteria* phylum. Considering the potentially high metabolic cost of maintaining such complex gene clusters in MTB, the wide observed MGC distribution across different lineages suggests that magnetosomal biogenesis and magnetotaxis must confer selective advantages on these organisms.

**Figure 4.**
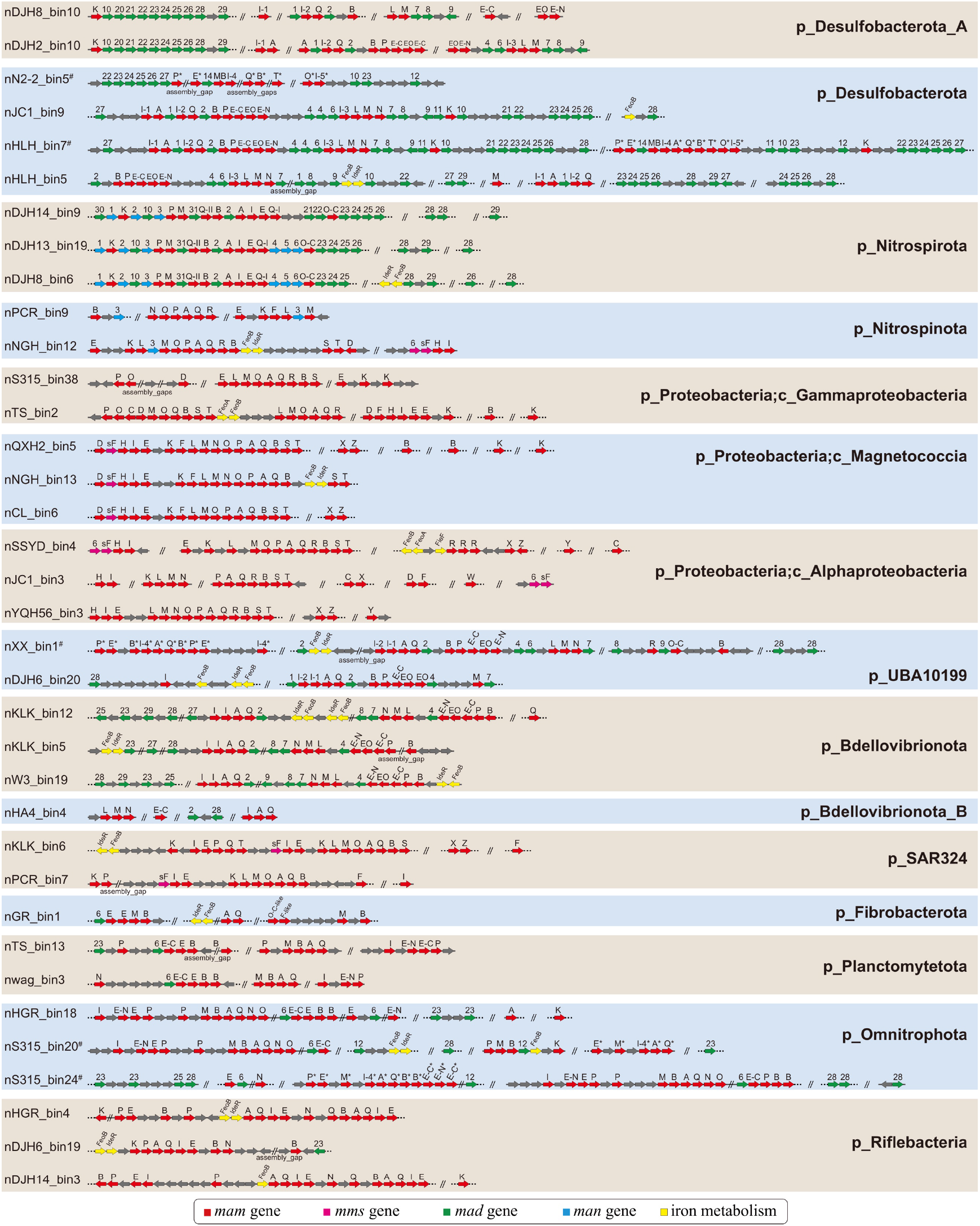
Representative magnetosome gene clusters (MGCs) from distinct MTB lineages recovered in this study. Genomes containing Fe_3_S_4_-type MGCs are highlighted with # and putative Fe_3_S_4_-type magnetosome genes in MGCs are denoted by *.

Two types of magnetosomal mineral crystals have been identified to date: magnetite (Fe_3_O_4_) and greigite (Fe_3_S_4_). Some MTB biomineralize both minerals within the same cell ^66,67^. Before this study, Fe_3_O_4_-type MGCs had been identified in all MTB lineages except for the *Latescibacterota* and *Planctomycetota* ^17^, while Fe_3_S_4_-type MGCs were only found in the *Desulfobacterota* (e.g., *Candidatus* Magnetoglobus multicellularis ^68^ and *Candidatus* Desulfamplus magnetomortis strain BW-1 ^67^), *Latescibacterota* ^36^, and *Planctomycetota* ^17^ phyla. Here, most of the identified MGCs contain Fe_3_O_4_-type magnetosome genes (including the first identification of Fe_3_O_4_-type MGCs in the *Planctomycetota* phylum) (Figure 4). Only a small fraction (nER2_bin1, nHLH_bin7, nN2-2_bin5, nS315_bin9, nS315_bin20, nS315_bin24, nTS_bin4, and nXX_bin1) harbor putative Fe_3_S_4_-type or both Fe_3_O_4_- and Fe_3_S_4_-type magnetosome genes, which suggests that Fe_3_O_4_-producing MTB are distributed more widely than Fe_3_S_4_-producing MTB in present-day habitats. The coexistence of Fe_3_O_4_- and Fe_3_S_4_-type MGCs in the same genomes of UBA10199 (nXX_bin1) and *Omnitrophota* (nS315_bin20 and nS315_bin24) phyla has never been reported previously, which not only indicates an unexpected phylogenetic diversity of Fe_3_S_4_-producing MTB, but, more importantly, challenges the traditional hypothesis regarding the origin of Fe_3_S_4_-producing MTB (discussed below). Thus, this study reveals that MTB in the domain *Bacteria* contain many more MGCs than anticipated, which captures a more complete picture of the genomic diversity of MTB. The diverse MGCs reconstructed here also represent a promising new gene resource for magnetic bio-nanoparticles that can be used to modify magnetosomal biogenesis pathways in MTB or even build new pathways in non-MTB^55^.

### Biogenesis of a magnetic organelle in the last Bacterial common ancestor (LBCA)?

A group of nine genes (*mamABEIKMOPQ*) was identified previously as the core magnetosome gene set shared by both Fe_3_O_4_- and Fe_3_S_4_-producing MTB ^61^. The products of these genes are thought to have important functions in magnetosomal biomineralization and for magnetosome chain construction. To trace the evolutionary history of magnetosomal biogenesis, we first performed a comparative genome analysis of 83 representative high-quality MTB genomes (with >90% completeness and <5% contamination) to identify core magnetosome genes that are defined such that >90% of input genomes (i.e., ≥74 genomes) must contain these genes, which allows for missing or fragmented genes due to the incomplete nature of draft genomes. Six magnetosome genes (*mamABIKMQ*) meet these criteria. Among these genes, proteins encoded by *mamBIMQ* are identified to be essential for magnetosomal biogenesis in *Magnetospirillum* strains; deletion of these genes results in non-magnetotactic mutants ^5,56,58^. Although *mamA* and *mamK* are not essential for magnetic mineral formation in *Magnetospirillum* species based on previous studies, the proteins they encoded are both involved in fine-tuning the magnetic dipole moment of the cell and thus magnetotaxis: MamA is responsible for magnetosome membrane assembly and MamK is involved in biomineral chain formation ^69–72^.

We then examined the phylogeny of magnetosome proteins MamABKMQ across available MTB genomes (Figure 5), with the exception of MamI due to its <60 aa positions after alignment trimming. The resulting trees are congruent overall with the genome-based phylogeny in that they are consistent with monophyly of the major phyla. This provides good evidence that the current MGC distribution across major Bacterial phyla is not due to extensive recent HGT events, but that it is due mainly to vertical inheritance. Coupled with the sharing of a common set of magnetosome genes, this strengthens the scenario of a single MGC emergence with later multiple independent losses during Bacterial diversification. However, a few discrepancies must be noted, such as separation between the UBA10199 phylum of nDJH6_bin20 and nXX_bin1 (Fe_3_O_4_-type), separate branching of nPCR_bin9 from the other *Nitrospinota* lineages, clustering of the Fe_3_O_4_-type *Planctomycetota* and *Omnitrophota* phyla, and clustering of *Desulfobacterota*_A and *Desulfobacterota* phyla in most protein trees, and clustering of *Nitrospinota* (except nPCR_bin9) and SAR324 within the *Proteobacteria* in trees of MamB and MamM (Figure 5 and Supplementary Figures 2 to 6). These discrepancies could have resulted from ancient HGT events. Alternatively, considering that only limited MTB genomes are available for the UBA10199, SAR324, *Nitrospinota*, and *Planctomycetota* phyla, these groupings could also be an artefact of tree reconstruction ^73^. Additional MTB genomes from these phyla will help to differentiate between these possibilities.

**Figure 5.**
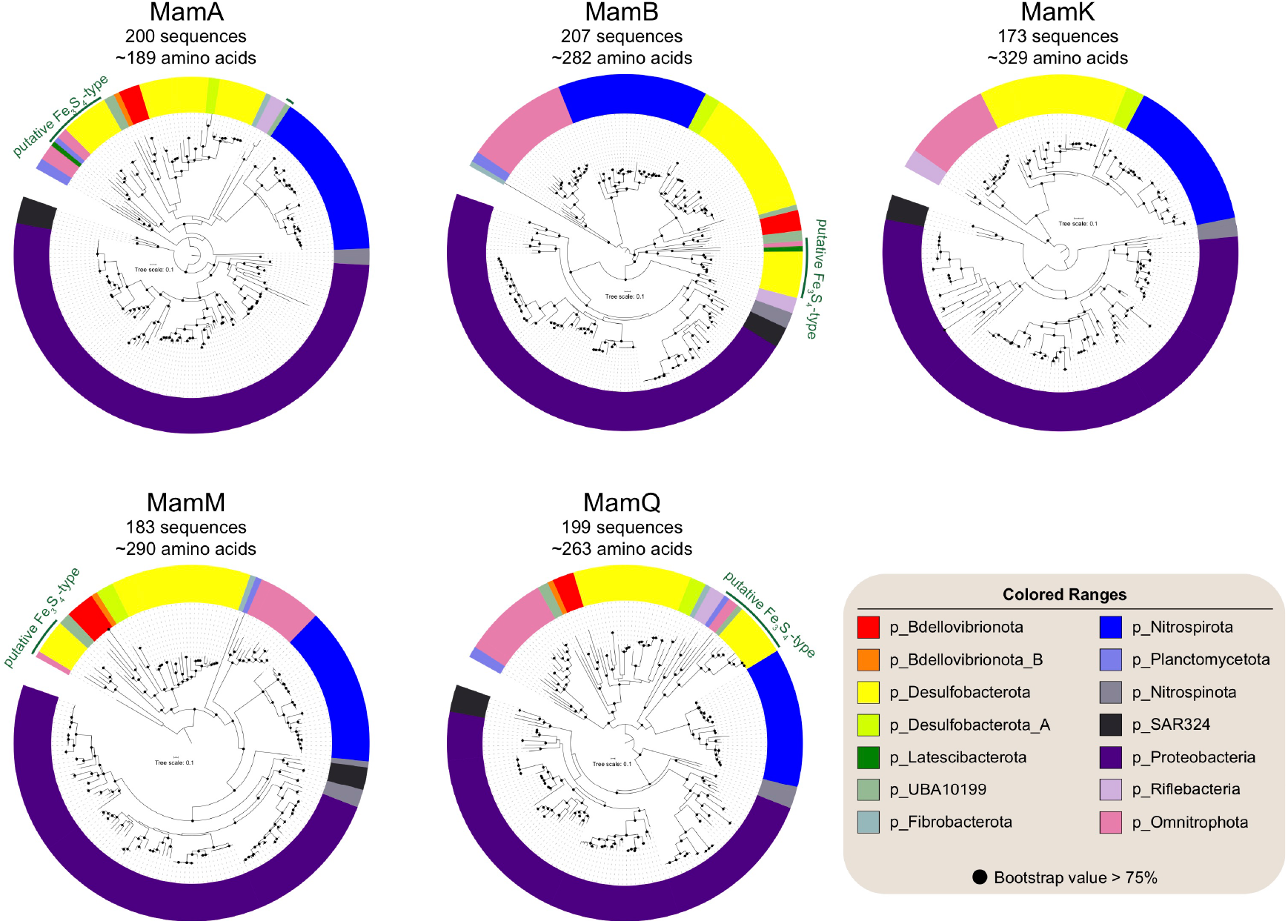
Maximum-likelihood trees of core magnetosome proteins. Trees were inferred using the MamABKMQ found in available MTB genomes. Complete trees are shown in Supplementary Figures 2 to 6.

Initial discovery of both Fe_3_O_4_- and Fe_3_S_4_-type MGCs within the same genome in the *Desulfobacterota* phylum (*Deltaproteobacteria* class in the NCBI taxonomy) led to the proposal that genes for Fe_3_S_4_ biomineralization in magnetosomes originated in this lineage ^15^. However, our finding of the coexistence of Fe_3_O_4_- and Fe_3_S_4_-type MGCs in the genomes of UBA10199 and *Omnitrophota* phyla weakens this hypothesis. Fe_3_S_4_- type magnetosome proteins form a monophyletic group in all MamABMQ trees with only the exception of Fe_3_S_4_-type MamA of nXX_bin1 that is clustered with *Nitrospirota* Fe_3_O_4_-type counterparts with low bootstrap support (<75%, Supplementary Figure 2). The structures of Fe_3_S_4_-type protein trees generally represent their phylum-level taxonomic phylogenies (Figure 5 and Supplementary Figures 2 to 6), which suggests that all Fe_3_S_4_-type magnetosome proteins likely shared a common ancestor and evolved separately in each phylum. These findings suggest that Fe_3_S_4_-type MGCs arose before the common ancestor of the Fe_3_S_4_-MTB containing phyla of *Desulfobacterota*, *Latescibacterota*, *Planctomycetota*, UBA10199, and *Omnitrophota*. The Fe_3_S_4_-type protein cluster is not associated robustly with any other Fe_3_O_4_-producing MTB lineages in protein trees, which precludes determination of the evolutionary origin of Fe_3_S_4_- producing MTB. Considering that Fe_3_O_4_-type MGCs are the most widespread, with identification in all major MTB lineages, it is likely that the Fe_3_O_4_-types are the ancestral form of magnetosome biosynthesis and that Fe_3_S_4_-type MGCs arose from Fe_3_O_4_-type MGCs through gene cluster duplication and divergence ^15^. However, both Fe_3_O_4_- and Fe_3_S_4_-type MGCs originating from an ancient unknown MGC type (Fe_3_S_4_ or other iron-containing biominerals) is also plausible ^37^.

Magnetosomal biogenesis is now considered to have an ancient origin ^15,37,41,74^. Discovery in this study of divergent MGCs across various Bacterial phyla expands significantly the database of MTB genomes and allows development of a more comprehensive scenario for the origin of magnetosomal biogenesis. The phylogenies of core magnetosome proteins (Figure 5) strengthens the notion of an ancient MGC origin, which then spread through a combination of vertical inheritance over geological times followed by multiple independent losses, potential HGT events, and gene/cluster duplications. This should have occurred from a more basal ancestor than previously thought prior to the divergence of all 14 known MTB-containing phyla (Figure 3). These phyla are scattered within the Bacterial tree of life and their common ancestor can be traced to near the base of the tree (Supplementary Figure 7), which indicates parsimoniously that close descendants of LBCA or the LBCA itself may have already contained ancestors of magnetosome genes and were, thus, capable of biomineralizing primitive magnetosomes.

Potential biogenesis of the magnetosome organelle in the LBCA has two major implications. First, the LBCA could have already possessed a relatively complex subcellular organization and, thus, was not as “primitive” as is usually imagined. Early organisms on Earth are described typically as simple organisms that lacked complex subcellular structures; however, a conceptual model of a complex LBCA or even a complex last universal common ancestor (LUCA) has been proposed based on the presence of eukaryote-like features present in some members of the *Planctomycetes* ^75,76^. Our results imply the formation of a magnetosome organelle in the LBCA, which supports the idea of a relatively complex LBCA. On early Earth the lack of a protective ozone layer resulted in higher harmful ultraviolet radiation than the present-day Earth, which would have been one of major challenges for life in the surface and shallow-water conditions ^77^. The intrinsic enzyme-like properties of magnetosome iron nanoparticles ^78,79^ and their potentially enhanced enzyme-like stability under a wide range of temperatures and pH ^80^ might have helped life to cope with environmental stresses on early Earth (e.g., detoxification of ultraviolet radiation and free-iron-generated ROS ^12^). No MGC has yet been found in the *Archaea*, which suggests that magnetosomal biogenesis evolved after the divergence of *Bacteria* and *Archaea*. Second, a magnetosome-forming LBCA implies that this feature may have been inherited by a taxonomically wide group of Bacterial phyla, while other phyla lost this trait during evolution. Thus, we argue that many MTB-containing lineages await discovery. Future discovery of additional MTB affiliated within other Bacterial phyla, especially those near the base of the Bacterial tree ^81,82^, will help to understand and constrain the evolutionary origin of the magnetosome organelle.

In summary, we reconstruct 168 MTB genomes here, which expand substantially the genomic representation of MTB and indicate a much more widespread distribution of magnetosome organelle biogenesis across the domain *Bacteria*. Analysis of core magnetosome proteins in the largest available taxonomic representation strengthens the notion of an ancient origin for magnetosome organelle biogenesis, which may date back to the LBCA. Genomes from this study will enable a better understanding of the biology and biomineralization of MTB and offer clues to assist in cultivation of uncultured MTB from different lineages.

## Methods

### Sample collection

A total of 53 sediment and soil samples were collected from a wide range of natural environments across China and Australia (Figure 2a), including 13 sediment samples that have been previously described ^37^ (for details see Supplementary Table 1). Each sample was examined for the presence of living MTB by light microscopy using the hanging-drop method ^83^. MTB cells from sediments and peatland soils were enriched magnetically using a “MTB trap” and enriched MTB cells were then subjected to metagenomic analyses, the detailed procedures for which are described elsewhere ^37,41,84^.

### Metagenome assembly, population genome binning, and comparative genomic analyses

Metagenomic DNA from each location was sequenced on Illumina HiSeq 2000, 2500, or 4000 platforms. Sequencing data were processed through a single-sample assembly and binning strategy using a MetaWRAP pipeline ^85^. The individual metagenomic datasets were assembled separately *de novo* using metaSPAdes (version 3.13.0) ^86^ with default parameters. Assembled scaffolds ≥2000 bp were binned separately using MetaBAT2 (version 2.12.1) ^87^, MaxBin2 (version 2.2.4) ^88^, and CONCOCT ^89^. Results of three binning methods for each sample were refined using MetaWRAP’s Bin_refinement and Reassemble_bins ^85^. Genome completeness and contamination were estimated with CheckM ^90^ using the ‘lineage_wf’ workflow. Only genomes with an estimated quality of >50 (defined as completeness − 5 × contamination) ^50^ were retained. Statistics for each genome were obtained using QUAST (version 4.2) ^91^, including the genome length, number of scaffolds, largest scaffold, GC content, and N50. Resultant genomes were annotated using Prokka (version 1.11) ^92^. Reconstructed genomes were checked manually for the presence of MGCs, followed by extensive manual verification of candidate magnetosome genes using NCBI PSI-BLAST ^93^. Notably, genome sequences with >99% ANI of previously reconstructed MTB genomes ^37^ from the same samples were recovered here using a different approach, which emphasizes the reproducibility of different genome-resolved metagenomic approaches. Taxonomic annotation of all acquired MTB genomes was performed using the Genome Taxonomy Database Toolkit GTDB-Tk ^53^ (version 0.3.2, database Release 04-RS89) with the ‘classify_wf’ function and default parameters. The genomes reconstructed here were combined with published MTB genomes (Supplementary Table 3). These genomes were dereplicated at 95% ANI for species ^51^ delineation using dRep ^94^ with ‘-sa 0.95’.

To identify the core magnetosome gene set shared by MTB in their respective genomes, only complete genomes and those with >90% completeness and <5% contamination are considered. These genomes were dereplicated using dRep ^94^ with ‘- sa 0.99’ for dereplication at 99% ANI, and finally 83 high-quality representative MTB genomes were selected. Most are draft genomes, so we define a core magnetosome protein as being present in >90% of the input genomes (i.e., at least 74 of 83 genomes) to minimize exclusion of potential core proteins due to the incomplete nature of draft genomes. Core proteins were calculated with COGtriangles ^95^ using GET_HOMOLOGUES ^96^ and were checked manually for magnetosome proteins. MamABIKMQ protein sequences were then searched with hidden Markov Models ^97^ (HMM) across known MTB genomes.

### Phylogenetic analyses

The phylogenetic tree composed of MTB genomes and those of relatively closely related non-MTB was inferred from 120 concatenated Bacterial single-copy marker proteins ^50^. Maximum-likelihood phylogeny was calculated using IQ-TREE (version 1.6.9) ^98^ under the LG+I+G4 substitution model with 1000 ultrafast bootstraps. The genome tree was rooted with the genome from the candidate phyla radiation (accession number LCFW00000000). For each MamABKMQ protein, sequences were aligned using MAFFT (version 7.407) ^99^ in ‘auto’ mode and filtered using trimAL ^100^ with ‘-gappyout’ option. Maximum-likelihood phylogenetic protein trees were then constructed using IQ-TREE under the TEST option for best model selection with 1000 ultrafast bootstraps. Protein trees were rooted at the midpoint. All trees were visualized using FigTree version 1.4.2 (http://tree.bio.ed.ac.uk/software/figtree/) and Interactive Tree Of Life (iTOL) v4 ^101^. AnnoTree ^102^ was used for phylogenomic visualization of the distribution of MTB-containing phyla across the Bacterial tree of life at the taxonomic phylum level.

### Data availability

Genome sequences are deposited in the NCBI BioProject under PRJNA400260 (accession numbers pending).

## Acknowledgments

We thank Xianyu Huang from State Key Laboratory of Biogeology and Environmental Geology, China University of Geosciences (Wuhan) and Zhiqi Zhang from Shennongjia National Park Administration for their contributions to the collection of acidic peatland soil samples. We thank Li Liu, Jia Liu, Runjia Ji, Yan Chen, and Yuan Fang for assistance with fieldwork. We thank Patrick De Deckker for suggesting field sampling sites and David Gordon and Samantha Burn for granting access to laboratory and materials at the Australian National University that enabled MTB extraction after our Australian fieldwork. This work was supported by the National Natural Science Foundation of China (NSFC) Grants 41621004 and 41822704, the Youth Innovation Promotion Association of the Chinese Academy of Sciences, the Natural Environment Research Council Independent Research Fellowship NE/P017266/1, and the Australian Research Council Grant DP140104544.

## Author contributions

W.L. and Y.X.P. conceived and planned the study. W.L., W.S.Z., G.A.P., X.Z., A.P.R., and Y.X.P. performed field sampling. W.L. and W.S.Z. enriched environmental magnetotactic bacteria and analyzed the data. W.L. wrote the manuscript with input from G.A.P., Q.Y.Z., R.K., D.A.B., A.P.R., and Y.X.P All authors approved the final manuscript.

## Supplementary materials

**Supplementary Figure 1.** Maximum likelihood phylogenomic tree of MTB genomes and their close non-MTB relatives.

**Supplementary Figure 2.** Maximum-likelihood tree of magnetosome protein MamA.

**Supplementary Figure 3.** Maximum-likelihood tree of magnetosome protein MamB.

**Supplementary Figure 4.** Maximum-likelihood tree of magnetosome protein MamK.

**Supplementary Figure 5.** Maximum-likelihood tree of magnetosome protein MamM.

**Supplementary Figure 6.** Maximum-likelihood tree of magnetosome protein MamQ.

**Supplementary Figure 7.** Phylogenetic distribution of MTB-containing phyla across the Bacterial tree of life. The phylum level Bacterial tree of life with MTB-containing phyla highlighted in blue. The Bacterial tree was made using the AnnoTree server.

**Supplementary Table 1.** Summary of sampled sites.

**Supplementary Table 2.** General characteristics of the 168 MTB genomes reported in this study. Genome completeness and contamination were estimated using CheckM and genome statistics were obtained using QUAST (version 4.2). Genome quality was defined as (completeness - 5 × contamination).

**Supplementary Table 3.** Previously published MTB genomes included in this study.

