## Supplementary for "Expanding magnetic organelle biogenesis in the domain *Bacteria*"

**Supplementary Figure 1.** Maximum likelihood phylogenomic tree of MTB genomes and their close non-MTB relatives.

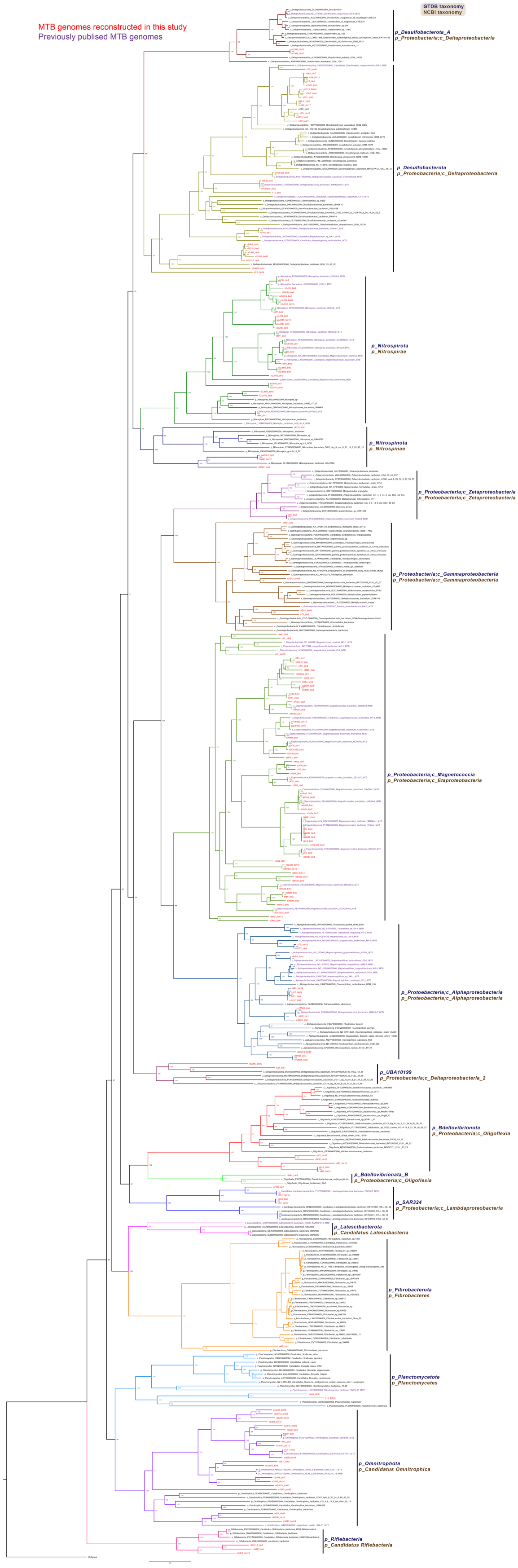

**Supplementary Figure 2.** Maximum-likelihood tree of magnetosome protein MamA.

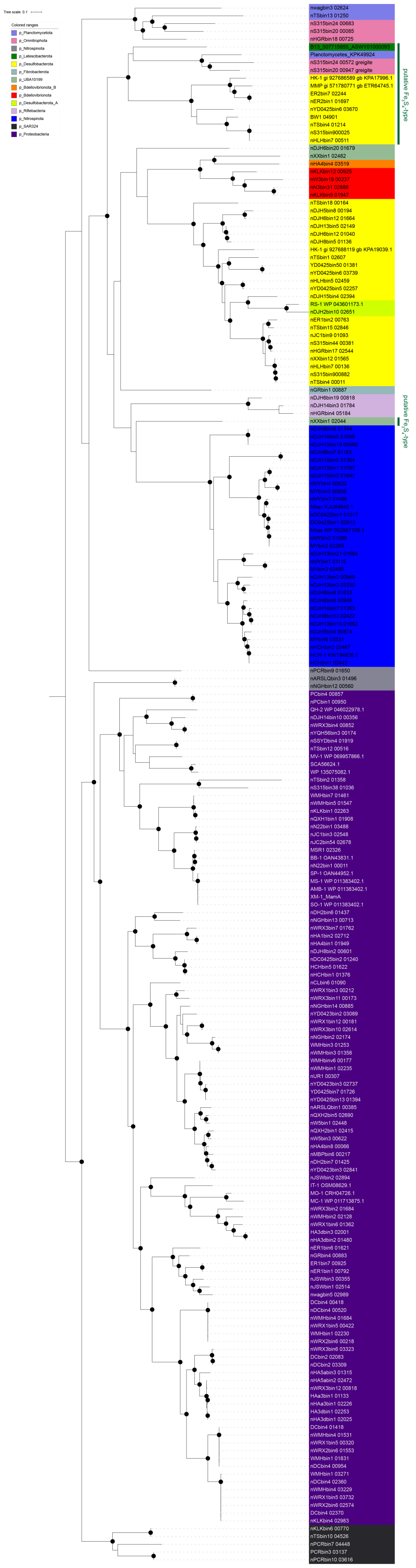

**Supplementary Figure 3.** Maximum-likelihood tree of magnetosome protein MamB.

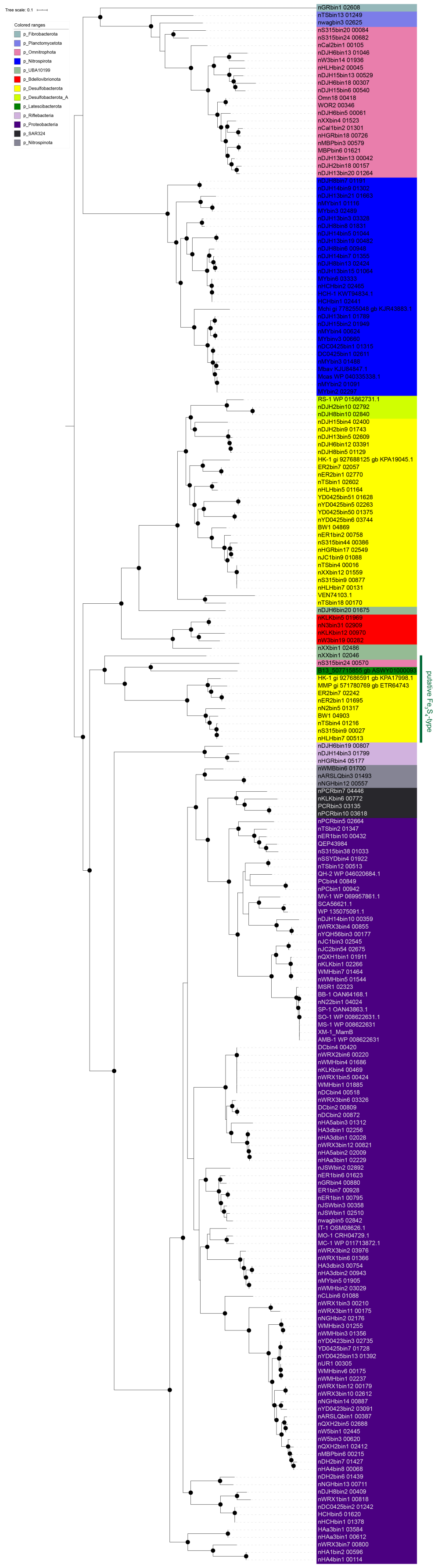

**Supplementary Figure 4.** Maximum-likelihood tree of magnetosome protein MamK.

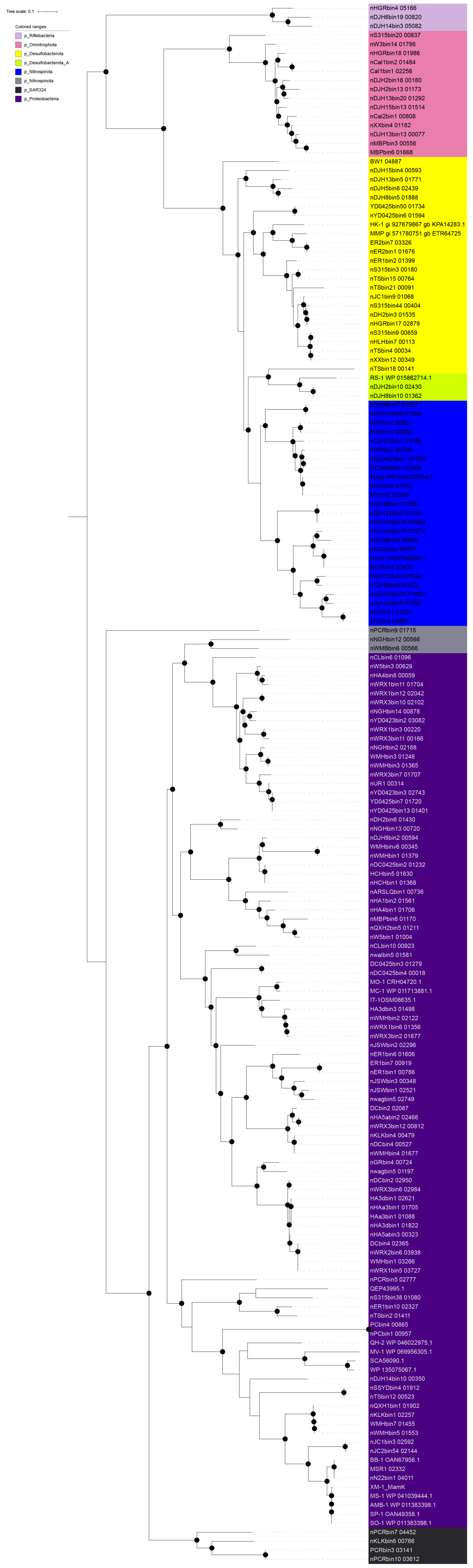

**Supplementary Figure 5.** Maximum-likelihood tree of magnetosome protein MamM.

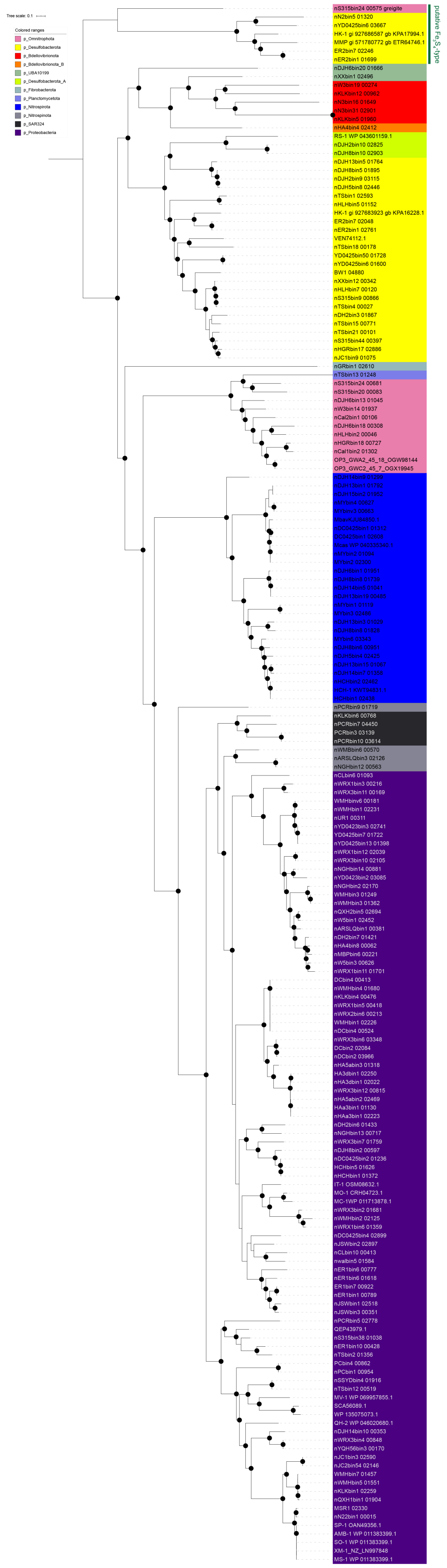

**Supplementary Figure 6.** Maximum-likelihood tree of magnetosome protein MamQ.

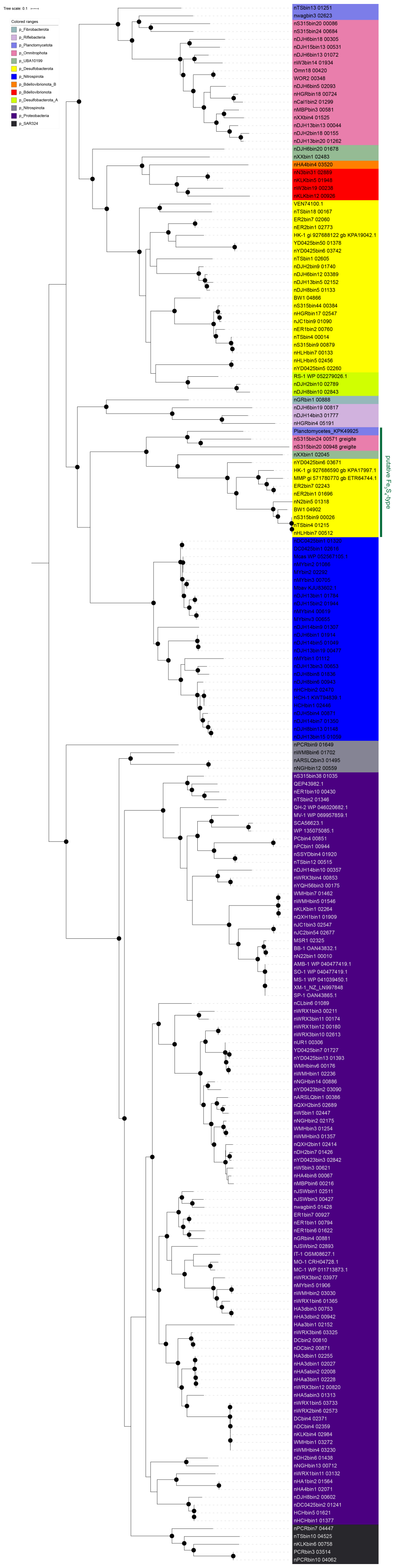

**Supplementary Figure 7.** Phylogenetic distribution of MTB-containing phyla across the Bacterial tree of life. The phylum level Bacterial tree of life with MTB-containing phyla highlighted in blue. The Bacterial tree was made using the AnnoTree server.

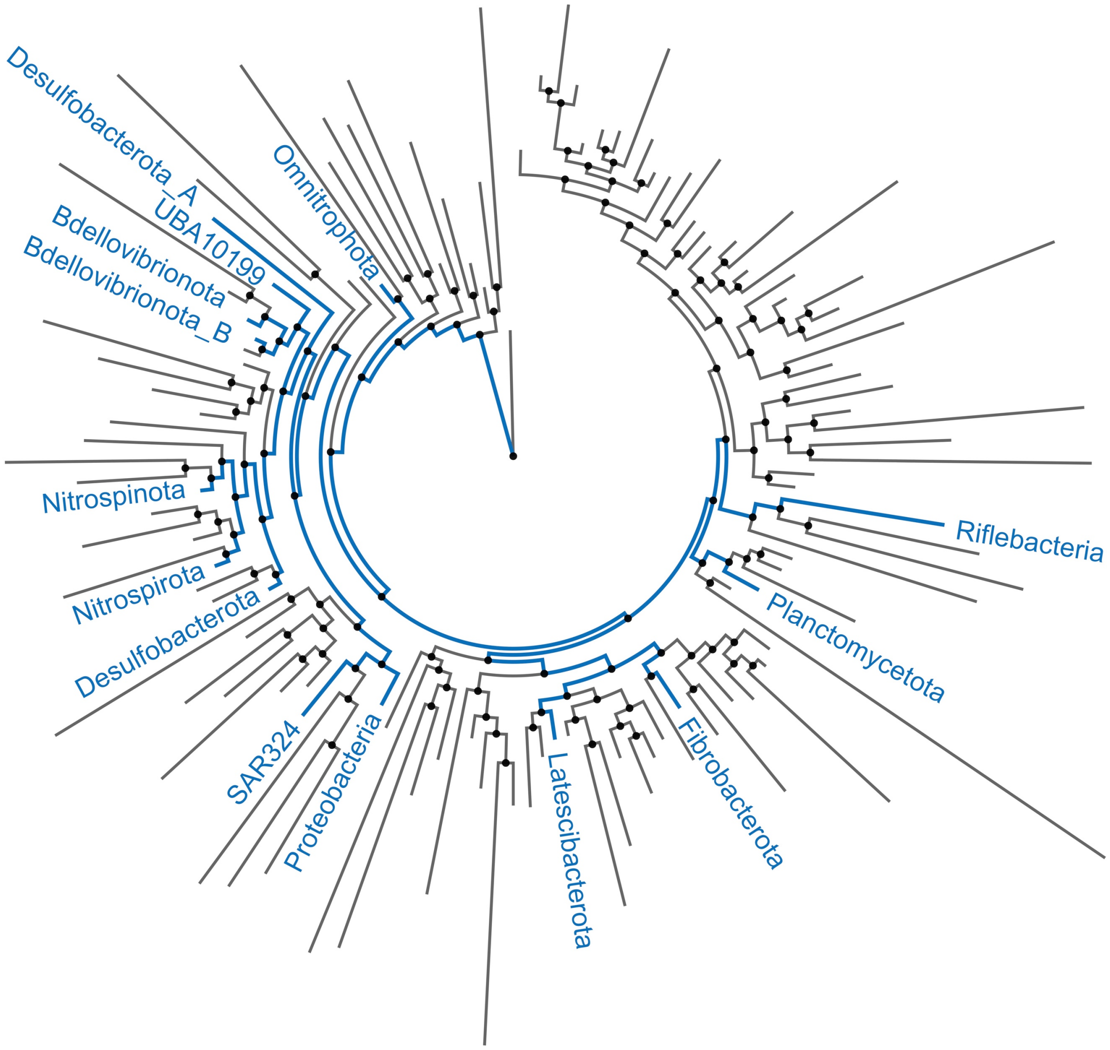

**Supplementary Table 1. Summary of sampled sites.**

| Sample ID | Sample location | Environment | Latitude (°) | Longitude (°E) | Salinity (ppt) | pH | Reference |
|---|---|---|---|---|---|---|---|
| ARSLQ | Innermogolia, China | Freshwater/Marginal | 47.19788 | 119.92877 | 0.3 | 6.8 | This study |
| Cal1 | Lake Catani, Australia | Freshwater/Marginal | -36.73426 | 146.81119 | <0.1 | / | Lin et al., 2018* |
| Cal2 | Lake Catani, Australia | Freshwater/Marginal | -36.73274 | 146.81226 | <0.1 | / | This study |
| CL | Corunna Lake, Australia | Freshwater/Marginal | -38.23694 | 144.42936 | 0.6 | / | This study |
| DC | Lake Dianchi, China | Freshwater/Marginal | 24.908133 | 102.74605 | 0.9 | 7.6 | Lin et al., 2018 |
| DC0425 | Lake Dianchi, China | Freshwater/Marginal | 24.908133 | 102.74605 | 0.9 | 7.6 | This study |
| DH2 | Lake East, China | Freshwater/Marginal | 30.555833 | 114.411389 | / | / | This study |
| DJH13 | Peatland, Dajiu Lake, China | Acidic Peatland | 31.483672 | 110.000694 | <0.1 | 5.1 | This study |
| DJH14 | Peatland, Dajiu Lake, China | Acidic Peatland | 31.483503 | 110.000844 | <0.1 | 4.9 | This study |
| DJH15 | Peatland, Dajiu Lake, China | Acidic Peatland | 31.485181 | 110.001483 | <0.1 | 5.7 | This study |
| DJH2 | Peatland, Dajiu Lake, China | Acidic Peatland | 31.49194 | 109.99472 | <0.1 | 4.9 | This study |
| DJH5 | Peatland, Dajiu Lake, China | Acidic Peatland | 31.490422 | 109.995886 | 0.1 | 4.4 | This study |
| DJH6 | Peatland, Dajiu Lake, China | Acidic Peatland | 31.49055 | 109.99611 | 0.2 | 4.9 | This study |
| DJH8 | Peatland, Dajiu Lake, China | Acidic Peatland | 31.48944 | 109.99638 | <0.1 | 4.3 | This study |
| ER1 | Erskine River, Australia | Brackish/Saline/Marine | -38.5338 | 143.97832 | 3.2 | / | Lin et al., 2018 |
| ER2 | Erskine River, Australia | Brackish/Saline/Marine | -38.5359 | 143.97478 | 1.9 | / | Lin et al., 2018 |
| GR | Gellibrand River, Australia | Brackish/Saline/Marine | -38.69715 | 143.15477 | 1.3 | / | This study |
| HA1 | Pond, Hongan, China | Freshwater/Marginal | 31.295 | 114.916944 | 0.4 | 7.3 | This study |
| HA3d | Pond, Hongan, China | Freshwater/Marginal | 31.173819 | 114.547642 | 0.2 | 5.2 | Lin et al., 2018 |
| HA4 | Pond, Hongan, China | Freshwater/Marginal | 31.292222 | 114.921667 | 0.3 | 7.3 | This study |
| HA5a | Pond, Hongan, China | Freshwater/Marginal | 31.290556 | 114.913333 | 0.4 | 7.5 | This study |
| HAa3 | Rice field, Hongan, China | Freshwater/Marginal | 31.174289 | 114.543941 | <0.1 | 5.2 | Lin et al., 2018 |
| HCH | Xi'an city moat, China | Freshwater/Marginal | 34.25287 | 108.92187 | 0.2 | 7.5 | Lin et al., 2018 |

| | | | | | | | |
|---|---|---|---|---|---|---|---|
| HGR | Honggaori Lake, Inner Mogolia, China | Freshwater/Marginal | 41.573047 | 108.313566 | 0.4 | 8.0 | This study |
| HLH | Hulun Lake, Inner Mogolia, China | Freshwater/Marginal | 49.06944 | 117.75694 | 0.7 | 8.6 | This study |
| JC1 | Pond, Inner Mogolia, China | Freshwater/Marginal | 41.687541 | 108.393862 | 0.3 | 8.2 | This study |
| JC2 | Pond, Inner Mogolia, China | Freshwater/Marginal | 41.686682 | 108.39418 | 0.3 | 8.2 | This study |
| JSW | Jinshawan, Yingkou, China | Brackish/Saline/Marine | 40.223972 | 122.083844 | / | / | This study |
| KLK | Keluke Lake, Qinghai, China | Brackish/Saline/Marine | 37.283722 | 96.859816 | 4.2 | 7.9 | This study |
| MBP | Mount Beauty Pondage, Australia | Freshwater/Marginal | -36.73874 | 147.16443 | <0.1 | / | Lin et al., 2018 |
| MY | Lake Miyun, China | Freshwater/Marginal | 40.48874 | 117.00714 | 0.2 | 7.5 | Lin et al., 2018 |
| N2-2 | Naritu, Inner Mogolia, China | Freshwater/Marginal | 41.549487 | 109.05952 | 0.3 | 7.5 | This study |
| N3 | Naritu, Inner Mogolia, China | Freshwater/Marginal | 41.549487 | 109.05952 | 0.8 | 7.8 | This study |
| NGH | Nuogan Lake, Inner Mogolia, China | Freshwater/Marginal | 47.9275 | 119.54638 | 0.3 | 7.4 | This study |
| PC | Painkalac Creek, Australia | Brackish/Saline/Marine | -38.46575 | 144.09288 | 21.7 | / | Lin et al., 2018 |
| PCR | Punkally Creek, Australia | Brackish/Saline/Marine | -36.23413 | 150.06798 | 33.9 | / | Lin et al., 2018 |
| QXH1 | Qixian Lake, Inner Mogolia, China | Freshwater/Marginal | 47.74777 | 119.33694 | 0.7 | 8.4 | This study |
| QXH2 | Qixian Lake, Inner Mogolia, China | Freshwater/Marginal | 47.69541 | 119.30422 | 0.3 | 8.5 | This study |
| S315 | Lake, Qinghai, China | Brackish/Saline/Marine | 38.229771 | 90.697681 | 4.3 | 8.5 | This study |
| SSYD | Shuishangyadan, Qinghai, China | Brackish/Saline/Marine | 37.624414 | 93.733812 | 15.3 | 7.6 | This study |
| TS | Tuosu Lake, Qinghai, China | Brackish/Saline/Marine | 37.197953 | 96.868083 | 3.8 | 7.8 | This study |
| W3 | Wulantaolegai, Inner Mogolia, China | Freshwater/Marginal | 41.564702 | 109.081861 | 0.4 | 8.1 | This study |
| W5 | Wulantaolegai, Inner Mogolia, China | Freshwater/Marginal | 41.564702 | 109.081861 | 0.4 | 8.1 | This study |
| wag | Waganaga Inlet, Australia | Brackish/Saline/Marine | -36.21831 | 150.12168 | 36.5 | / | This study |
| wal | Wallaga Lake, Australia | Brackish/Saline/Marine | -36.36013 | 150.07384 | 37.0 | / | This study |
| WMH | Lake Weiming, China | Freshwater/Marginal | 39.993142 | 116.30256 | 0.3 | 7.5 | Lin et al., 2018 |
| WRX1 | Weierxun river, Inner Mogolia, China | Freshwater/Marginal | 48.23222 | 117.62305 | 0.2 | 8.0 | This study |
| WRX2 | Weierxun river, Inner Mogolia, China | Freshwater/Marginal | 48.23111 | 117.62277 | 0.3 | 7.9 | This study |

| | | | | | | | |
|---|---|---|---|---|---|---|---|
| WRX3 | Weierxun river, Inner Mogolia, China | Freshwater/Marginal | 48.455 | 117.63305 | 0.2 | 7.9 | This study |
| XX | Aiken, creek, Qinghai, China | Freshwater/Marginal | 38.170074 | 90.57854 | 0.8 | 7.7 | This study |
| YD0423 | Yuandadu Park, China | Freshwater/Marginal | 39.974732 | 116.368688 | 0.5 | 7.2 | This study |
| YD0425 | Yuandadu Park, China | Freshwater/Marginal | 39.974732 | 116.368688 | 0.5 | 7.2 | Lin et al., 2018 |
| YQH56 | Yanqi Lake, Beijing, China | Freshwater/Marginal | 40.39833 | 116.67777 | / | / | This study |

*Lin W, Zhang W, Zhao X, et al. Genomic expansion of magnetotactic bacteria reveals an early common origin of magnetotaxis with lineage-specific evolution. ISME J, 2018, 12:1508–1519.

**Supplementary Table 2. General characteristics of the 168 MTB genomes reported in this study.** Genome completeness and contamination were estimated using CheckM and genome statistics were obtained using QUAST (version 4.2). Genome quality was defined as (completeness - 5 × contamination).

| Genome ID | Completeness (%) | Contamination (%) | Quality | Number of scaffolds | Largest scaffold (bp) | Total length (bp) | GC (%) | N50 | GTDB classification |
|---|---|---|---|---|---|---|---|---|---|
| nARSLQ_bin1 | 70.17 | 0 | 70.17 | 161 | 188805 | 3310492 | 58.83 | 41237 | d__Bacteria;p__Proteobacteria;c__Magnetococcia;o__Magnetococcales;f__WMHbin3;g__WMHbin3;s__ |
| nARSLQ_bin3 | 76.98 | 0.05 | 76.73 | 440 | 43549 | 2637314 | 54.88 | 7942 | d__Bacteria;p__Nitrospinota;c__UBA7883;o__UBA7883;f__UBA7883;g__;s__ |
| nCal1_bin2 | 88.29 | 2.15 | 77.54 | 196 | 108977 | 2952830 | 49.66 | 25022 | d__Bacteria;p__Omnitrophota;c__koll11;o__UBA10015;f__GCA-002753745;g__GCA-2753745;s__GCA-2753745 sp002753745 |
| nCal2_bin1 | 83.95 | 3.23 | 67.8 | 176 | 47664 | 1746582 | 51.48 | 15478 | d__Bacteria;p__Omnitrophota;c__koll11;o__UBA10015;f__GCA-002753745;g__GCA-2753745;s__ |
| nCL_bin10 | 61.19 | 2.1 | 50.69 | 364 | 93268 | 3358134 | 55.62 | 18428 | d__Bacteria;p__Proteobacteria;c__Magnetococcia;o__Magnetococcales;f__;g__;s__ |
| nCL_bin6 | 86.9 | 2.94 | 72.2 | 117 | 324607 | 3646512 | 47.56 | 61959 | d__Bacteria;p__Proteobacteria;c__Magnetococcia;o__Magnetococcales;f__;g__;s__ |
| nDC_bin2 | 94.91 | 2.52 | 82.31 | 311 | 95666 | 4466229 | 51.87 | 24333 | d__Bacteria;p__Proteobacteria;c__Magnetococcia;o__Magnetococcales;f__UBA8363;g__UBA8363;s__UBA8363 sp002753615 |
| nDC_bin4 | 97.48 | 2.1 | 86.98 | 149 | 180801 | 4493558 | 54.13 | 47506 | d__Bacteria;p__Proteobacteria;c__Magnetococcia;o__Magnetococcales;f__UBA8363;g__UBA8363;s__UBA8363 sp002753735 |
| nDC0425_bin1 | 96.97 | 1.82 | 87.87 | 178 | 169563 | 4037616 | 49.09 | 35907 | d__Bacteria;p__Nitrospirota;c__Thermodesulfovibrionia;o__Thermodesulfovibrionales;f__Magnetobacteriaceae;g__Magnetobacterium;s__Magnetobacterium sp002753685 |
| nDC0425_bin2 | 71.32 | 1.79 | 62.37 | 364 | 59974 | 3037918 | 55.86 | 10610 | d__Bacteria;p__Proteobacteria;c__Magnetococcia;o__Magnetococcales;f__WMHbin3;g__HCHbin5;s__HCHbin5 sp002753505 |

| | | | | | | | | | |
|---|---|---|---|---|---|---|---|---|---|
| nDC0425_bin4 | 98.32 | 1.68 | 89.92 | 165 | 135657 | 3892739 | 65.39 | 49112 | d__Bacteria;p__Proteobacteria;c__Magnetococcia;o__Magnetococcales;f__DC0425bin3;g__DC0425bin3;s__DC0425bin3 sp002753665 |
| nDH2_bin3 | 75.5 | 1.29 | 69.05 | 407 | 39022 | 2540914 | 35.65 | 7726 | d__Bacteria;p__Desulfobacterota;c__Desulfobacteria;o__Desulfobacterales;f__Desulfobacteraceae;g__Desulfamplus;s__ |
| nDH2_bin6 | 80.77 | 2.1 | 70.27 | 414 | 83996 | 4543253 | 52.7 | 21120 | d__Bacteria;p__Proteobacteria;c__Magnetococcia;o__Magnetococcales;f__;g__;s__ |
| nDH2_bin7 | 54.48 | 0 | 54.48 | 74 | 184375 | 1952691 | 56.31 | 45589 | d__Bacteria;p__Proteobacteria;c__Magnetococcia;o__Magnetococcales;f__WMHbin3;g__WMHbin3;s__ |
| nDJH13_bin1 | 85.91 | 0.45 | 83.66 | 168 | 88709 | 3320021 | 49.07 | 29980 | d__Bacteria;p__Nitrospirota;c__Thermodesulfovibrionia;o__Thermodesulfovibrionales;f__Magnetobacteriaceae;g__Magnetobacterium;s__ |
| nDJH13_bin13 | 54.51 | 0 | 54.51 | 97 | 48009 | 1173800 | 54.15 | 15478 | d__Bacteria;p__Omnitrophota;c__koll11;o__UBA10015;f__GCA-002753745;g__GCA-2753745;s__ |
| nDJH13_bin15 | 66.54 | 0 | 66.54 | 121 | 58913 | 2034651 | 48.07 | 27741 | d__Bacteria;p__Nitrospirota;c__Thermodesulfovibrionia;o__Thermodesulfovibrionales;f__Magnetobacteriaceae;g__HCH-1;s__ |
| nDJH13_bin19 | 99.03 | 0.91 | 94.48 | 65 | 477006 | 3711429 | 41.57 | 125510 | d__Bacteria;p__Nitrospirota;c__Thermodesulfovibrionia;o__Thermodesulfovibrionales;f__Magnetobacteriaceae;g__;s__ |
| nDJH13_bin20 | 60.34 | 0 | 60.34 | 31 | 321724 | 2119943 | 42.8 | 179029 | d__Bacteria;p__Omnitrophota;c__koll11;o__UBA10015;f__GCA-002753745;g__;s__ |
| nDJH13_bin21 | 85.99 | 2.25 | 74.74 | 262 | 98345 | 3287229 | 47.98 | 20424 | d__Bacteria;p__Nitrospirota;c__Thermodesulfovibrionia;o__Thermodesulfovibrionales;f__UBA9935;g__GCA-2634385;s__ |
| nDJH13_bin3 | 77.94 | 1.82 | 68.84 | 220 | 106824 | 4006792 | 47.81 | 25191 | d__Bacteria;p__Nitrospirota;c__Thermodesulfovibrionia;o__Thermodesulfovibrionales;f__Magnetobacteriaceae;g__;s__ |
| nDJH13_bin5 | 56.21 | 0 | 56.21 | 619 | 51923 | 4587119 | 51.85 | 9499 | d__Bacteria;p__Desulfobacterota;c__Desulfobacteria;o__;f__;g__;s__ |
| nDJH14_bin10 | 73.86 | 1.33 | 67.21 | 435 | 52301 | 3326403 | 60.2 | 11257 | d__Bacteria;p__Proteobacteria;c__Alphaproteobacteria;o__Rhodospirillales_B;f__;g__;s__ |

| | | | | | | | | | |
|---|---|---|---|---|---|---|---|---|---|
| nDJH14_bin3 | 96 | 1.33 | 89.35 | 185 | 380595 | 8297085 | 40.61 | 109594 | d__Bacteria;p__Riflebacteria;c__UBA8953;o__UBA8953;f__UBA8953;g__;s__ |
| nDJH14_bin5 | 95.85 | 0.91 | 91.3 | 69 | 359807 | 3591102 | 41.5 | 83655 | d__Bacteria;p__Nitrospirota;c__Thermodesulfovibrionia;o__Thermodesulfovibrionales;f__Magnetobacteriaceae;g__;s__ |
| nDJH14_bin7 | 85.56 | 0.45 | 83.31 | 565 | 37924 | 4238609 | 44.94 | 10032 | d__Bacteria;p__Nitrospirota;c__Thermodesulfovibrionia;o__Thermodesulfovibrionales;f__Magnetobacteriaceae;g__HCH-1;s__ |
| nDJH14_bin9 | 95.95 | 0.91 | 91.4 | 75 | 256720 | 3130381 | 35.21 | 93402 | d__Bacteria;p__Nitrospirota;c__Thermodesulfovibrionia;o__Thermodesulfovibrionales;f__Magnetobacteriaceae;g__;s__ |
| nDJH15_bin13 | 83.33 | 1.08 | 77.93 | 111 | 187318 | 2101230 | 49.08 | 34545 | d__Bacteria;p__Omnitrophota;c__koll11;o__UBA10015;f__kpj58rc;g__;s__ |
| nDJH15_bin2 | 92.02 | 0.91 | 87.47 | 245 | 111317 | 3580076 | 49.18 | 21641 | d__Bacteria;p__Nitrospirota;c__Thermodesulfovibrionia;o__Thermodesulfovibrionales;f__Magnetobacteriaceae;g__Magnetobacterium;s__ |
| nDJH15_bin4 | 60.62 | 0.32 | 59.02 | 701 | 64630 | 4202781 | 51.2 | 7918 | d__Bacteria;p__Desulfobacterota;c__Desulfarculia;o__Adiutricales;f__;g__;s__ |
| nDJH15_bin6 | 79.69 | 2.15 | 68.94 | 47 | 304738 | 2379669 | 43.25 | 130162 | d__Bacteria;p__Omnitrophota;c__koll11;o__UBA10015;f__kpj58rc;g__;s__ |
| nDJH15_bin8 | 64.82 | 1.82 | 55.72 | 62 | 155282 | 2181452 | 48.65 | 52271 | d__Bacteria;p__Nitrospirota;c__Thermodesulfovibrionia;o__Thermodesulfovibrionales;f__UBA9935;g__GCA-2634385;s__ |
| nDJH2_bin10 | 97.32 | 0 | 97.32 | 158 | 125371 | 3372092 | 62.81 | 36054 | d__Bacteria;p__Desulfobacterota_A;c__Desulfovibrionia;o__Desulfovibrionales;f__Desulfovibrionaceae;g__;s__ |
| nDJH2_bin13 | 70.95 | 0 | 70.95 | 209 | 47316 | 1763776 | 42.08 | 11646 | d__Bacteria;p__Omnitrophota;c__koll11;o__UBA10015;f__GCA-002753745;g__;s__ |
| nDJH2_bin18 | 61.47 | 1.08 | 56.07 | 38 | 212303 | 1845990 | 44.09 | 102146 | d__Bacteria;p__Omnitrophota;c__koll11;o__UBA10015;f__GCA-002753745;g__;s__ |
| nDJH2_bin9 | 72.63 | 1.06 | 67.33 | 633 | 46331 | 4764272 | 51.72 | 10360 | d__Bacteria;p__Desulfobacterota;c__Desulfobaccia_A;o__RBG-13-43-22;f__;g__;s__ |

| | | | | | | | | | |
|---|---|---|---|---|---|---|---|---|---|
| nDJH5_bin4 | 69.64 | 0 | 69.64 | 344 | 40054 | 2542339 | 47.41 | 11135 | d__Bacteria;p__Nitrospirota;c__Thermodesulfovibrionia;o__Thermodesulfovibrionales;f__Magnetobacteriaceae;g__HCH-1;s__ |
| nDJH5_bin8 | 67.24 | 0.65 | 63.99 | 521 | 39776 | 3747650 | 51.53 | 8844 | d__Bacteria;p__Desulfobacterota;c__Desulfobaccia_A;o__RBG-13-43-22;f__;g__;s__ |
| nDJH6_bin1 | 95.17 | 1.97 | 85.32 | 318 | 88236 | 3469571 | 41.13 | 17243 | d__Bacteria;p__Nitrospirota;c__Thermodesulfovibrionia;o__Thermodesulfovibrionales;f__Magnetobacteriaceae;g__;s__ |
| nDJH6_bin12 | 66.54 | 0.97 | 61.69 | 569 | 48544 | 3952401 | 51.41 | 9135 | d__Bacteria;p__Desulfobacterota;c__Desulfobaccia_A;o__RBG-13-43-22;f__;g__;s__ |
| nDJH6_bin13 | 73.12 | 0.1 | 72.62 | 175 | 88575 | 1591332 | 47.17 | 11632 | d__Bacteria;p__Omnitrophota;c__koll11;o__UBA1560;f__Omnitrophaceae_A;g__;s__ |
| nDJH6_bin14 | 66.9 | 1.72 | 58.3 | 50 | 165984 | 1979993 | 37.49 | 86148 | d__Bacteria;p__Omnitrophota;c__koll11;o__UBA10015;f__kpj58rc;g__;s__ |
| nDJH6_bin18 | 76.15 | 3.23 | 60 | 187 | 113774 | 2288015 | 41.64 | 23939 | d__Bacteria;p__Omnitrophota;c__koll11;o__UBA10015;f__GCA-002753745;g__;s__ |
| nDJH6_bin19 | 77.53 | 1.69 | 69.08 | 185 | 268319 | 6314202 | 49.32 | 78585 | d__Bacteria;p__Riflebacteria;c__UBA8953;o__UBA8953;f__UBA8953;g__;s__ |
| nDJH6_bin20 | 94.84 | 0.83 | 90.69 | 64 | 162140 | 2614640 | 45.14 | 63680 | d__Bacteria;p__UBA10199;c__UBA10199;o__;f__;g__;s__ |
| nDJH6_bin28 | 65.52 | 0 | 65.52 | 58 | 231325 | 1814106 | 47.32 | 63386 | d__Bacteria;p__Omnitrophota;c__koll11;o__UBA10015;f__GCA-002753745;g__;s__ |
| nDJH6_bin5 | 82.8 | 3.23 | 66.65 | 37 | 269155 | 2393593 | 40.68 | 108715 | d__Bacteria;p__Omnitrophota;c__koll11;o__UBA10015;f__kpj58rc;g__;s__ |
| nDJH8_bin10 | 95.77 | 0 | 95.77 | 255 | 122233 | 3285386 | 62.71 | 21540 | d__Bacteria;p__Desulfobacterota_A;c__Desulfovibrionia;o__Desulfovibrionales;f__Desulfovibrionaceae;g__;s__ |
| nDJH8_bin13 | 71.27 | 1.82 | 62.17 | 190 | 68691 | 2611585 | 47.66 | 19254 | d__Bacteria;p__Nitrospirota;c__Thermodesulfovibrionia;o__Thermodesulfovibrionales;f__Magnetobacteriaceae;g__HCH-1;s__ |

| | | | | | | | | | |
|---|---|---|---|---|---|---|---|---|---|
| nDJH8_bin2 | 98.99 | 3.99 | 79.04 | 434 | 142806 | 4525783 | 57.45 | 14666 | d__Bacteria;p__Proteobacteria;c__Magnetococcia;o__Magnetococcales;f__WMHbin3;g__HCHbin5;s__ |
| nDJH8_bin5 | 81.25 | 0.65 | 78 | 527 | 64082 | 4058279 | 51.56 | 10433 | d__Bacteria;p__Desulfobacterota;c__Desulfobaccia_A;o__RBG-13-43-22;f__;g__;s__ |
| nDJH8_bin6 | 98.03 | 0.91 | 93.48 | 149 | 138495 | 3781477 | 46.6 | 45032 | d__Bacteria;p__Nitrospirota;c__Thermodesulfovibrionia;o__Thermodesulfovibrionales;f__Magnetobacteriaceae;g__HCH-1;s__ |
| nDJH8_bin7 | 55.23 | 0 | 55.23 | 354 | 42065 | 1913775 | 35.15 | 6641 | d__Bacteria;p__Nitrospirota;c__Thermodesulfovibrionia;o__Thermodesulfovibrionales;f__Magnetobacteriaceae;g__;s__ |
| nDJH8_bin8 | 98.58 | 0.91 | 94.03 | 111 | 217288 | 4187783 | 41.49 | 60346 | d__Bacteria;p__Nitrospirota;c__Thermodesulfovibrionia;o__Thermodesulfovibrionales;f__Magnetobacteriaceae;g__;s__ |
| nER1_bin1 | 99.11 | 1.68 | 90.71 | 47 | 309656 | 3664887 | 52.32 | 113822 | d__Bacteria;p__Proteobacteria;c__Magnetococcia;o__Magnetococcales;f__UBA8363;g__GCA-2753565;s__GCA-2753565 sp002753565 |
| nER1_bin10 | 68.81 | 0.62 | 65.71 | 485 | 29143 | 2595233 | 54.7 | 6203 | d__Bacteria;p__Proteobacteria;c__Gammaproteobacteria;o__Thiohalomonadales;f__Thiohalomonadaceae;g__;s__ |
| nER1_bin2 | 73.13 | 0.97 | 68.28 | 486 | 62816 | 2882814 | 42.66 | 7321 | d__Bacteria;p__Desulfobacterota;c__Desulfobacteria;o__Desulfobacterales;f__Desulfobacteraceae;g__Desulfamplus;s__ |
| nER1_bin6 | 73.79 | 1.72 | 65.19 | 102 | 225238 | 3093236 | 51.04 | 51978 | d__Bacteria;p__Proteobacteria;c__Magnetococcia;o__Magnetococcales;f__UBA8363;g__GCA-2753565;s__ |
| nER2_bin1 | 93.82 | 0 | 93.82 | 393 | 107480 | 6162735 | 38.94 | 22709 | d__Bacteria;p__Desulfobacterota;c__Desulfobacteria;o__Desulfobacterales;f__Magnetomoraceae;g__Magnetomorum;s__Magnetomorum sp002753725 |
| nGR_bin1 | 88.69 | 1.1 | 83.19 | 350 | 195322 | 5376203 | 40.73 | 27450 | d__Bacteria;p__Fibrobacterota;c__Fibrobacteria;o__UBA11236;f__;g__;s__ |
| nGR_bin4 | 61.47 | 1.68 | 53.07 | 466 | 32025 | 2868495 | 42 | 7505 | d__Bacteria;p__Proteobacteria;c__Magnetococcia;o__Magnetococcales;f__UBA8363;g__GCA-2753565;s__ |
| nHA1_bin2 | 76.53 | 3.36 | 59.73 | 270 | 89738 | 3028678 | 50.29 | 17143 | d__Bacteria;p__Proteobacteria;c__Magnetococcia;o__Magnetococcales;f__WMHbin3;g__WMHbin3;s__ |

| | | | | | | | | | |
|---|---|---|---|---|---|---|---|---|---|
| nHA3d_bin1 | 97.41 | 2.94 | 82.71 | 203 | 189792 | 4465495 | 52.89 | 43128 | d__Bacteria;p__Proteobacteria;c__Magnetococcia;o__Magnetococcales;f__UBA8363;g__UBA8363;s__UBA8363 sp002753515 |
| nHA3d_bin2 | 86.97 | 3.78 | 68.07 | 358 | 109622 | 3545572 | 61.56 | 17432 | d__Bacteria;p__Proteobacteria;c__Magnetococcia;o__Magnetococcales;f__DC0425bin3;g__HA3dbin3;s__HA3dbin3 sp002753495 |
| nHA4_bin1 | 56.19 | 0 | 56.19 | 414 | 30892 | 2438036 | 50.34 | 7694 | d__Bacteria;p__Proteobacteria;c__Magnetococcia;o__Magnetococcales;f__WMHbin3;g__WMHbin3;s__ |
| nHA4_bin4 | 63.54 | 0.93 | 58.89 | 942 | 61771 | 4285723 | 43.67 | 5119 | d__Bacteria;p__Bdellovibrionota_B;c__Oligoflexia;o__Oligoflexales;f__Oligoflexaceae;g__;s__ |
| nHA4_bin8 | 53.73 | 0.05 | 53.48 | 382 | 47826 | 2053852 | 58.19 | 6996 | d__Bacteria;p__Proteobacteria;c__Magnetococcia;o__Magnetococcales;f__WMHbin3;g__;s__ |
| nHA5a_bin2 | 84.03 | 4.62 | 60.93 | 229 | 71658 | 3237221 | 52.36 | 25023 | d__Bacteria;p__Proteobacteria;c__Magnetococcia;o__Magnetococcales;f__UBA8363;g__UBA8363;s__UBA8363 sp002753515 |
| nHA5a_bin3 | 70.28 | 1.68 | 61.88 | 426 | 77980 | 3385493 | 54.67 | 12490 | d__Bacteria;p__Proteobacteria;c__Magnetococcia;o__Magnetococcales;f__UBA8363;g__UBA8363;s__ |
| nHAa3_bin1 | 81.93 | 1.68 | 73.53 | 97 | 201489 | 3004780 | 53.21 | 84180 | d__Bacteria;p__Proteobacteria;c__Magnetococcia;o__Magnetococcales;f__UBA8363;g__UBA8363;s__UBA8363 sp002753515 |
| nHCH_bin1 | 94.79 | 2.1 | 84.29 | 295 | 185742 | 3805225 | 56.77 | 22140 | d__Bacteria;p__Proteobacteria;c__Magnetococcia;o__Magnetococcales;f__WMHbin3;g__HCHbin5;s__HCHbin5 sp002753505 |
| nHCH_bin2 | 98.18 | 0.91 | 93.63 | 99 | 185786 | 3701936 | 45.24 | 57400 | d__Bacteria;p__Nitrospirota;c__Thermodesulfovibrionia;o__Thermodesulfovibrionales;f__Magnetobacteriaceae;g__HCH-1;s__HCH-1 sp001541255 |
| nHGR_bin17 | 95.65 | 1.97 | 85.8 | 106 | 182570 | 3818984 | 37.82 | 58122 | d__Bacteria;p__Desulfobacterota;c__Desulfobacteria;o__Desulfobacterales;f__Desulfobacteraceae;g__Desulfamplus;s__ |
| nHGR_bin18 | 88.29 | 2.15 | 77.54 | 80 | 141577 | 2643610 | 51.11 | 53313 | d__Bacteria;p__Omnitrophota;c__koll11;o__UBA10015;f__GCA-002753745;g__GCA-2753745;s__ |
| nHGR_bin4 | 96 | 0 | 96 | 54 | 713530 | 7047312 | 40.28 | 283247 | d__Bacteria;p__Riflebacteria;c__UBA8953;o__UBA8953;f__UBA8953;g__;s__ |

| | | | | | | | | | |
|---|---|---|---|---|---|---|---|---|---|
| nHLH_bin2 | 73.59 | 2.21 | 62.54 | 210 | 112777 | 2207557 | 42.78 | 15244 | d__Bacteria;p__Omnitrophota;c__koll11;o__UBA10015;f__kpj58rc;g__;s__ |
| nHLH_bin5 | 82.29 | 0.81 | 78.24 | 421 | 75920 | 4583064 | 32.52 | 15842 | d__Bacteria;p__Desulfobacterota;c__Desulfobacteria;o__Desulfobacterales;f__YD0425bin51;g__YD0425bin51;s__YD0425bin51 sp002753225 |
| nHLH_bin7 | 95.67 | 1.29 | 89.22 | 147 | 86214 | 3017047 | 37.75 | 34217 | d__Bacteria;p__Desulfobacterota;c__Desulfobacteria;o__Desulfobacterales;f__Desulfobacteraceae;g__Desulfamplus;s__ |
| nJC1_bin3 | 84.69 | 0.5 | 82.19 | 698 | 34368 | 3955744 | 67.27 | 6954 | d__Bacteria;p__Proteobacteria;c__Alphaproteobacteria;o__Rhodospirillales_B;f__Magnetospirillaceae;g__;s__ |
| nJC1_bin9 | 99.68 | 1.61 | 91.63 | 82 | 545051 | 4348730 | 35.99 | 138975 | d__Bacteria;p__Desulfobacterota;c__Desulfobacteria;o__Desulfobacterales;f__Desulfobacteraceae;g__Desulfamplus;s__ |
| nJC2_bin54 | 81.55 | 0.25 | 80.3 | 694 | 58916 | 3885447 | 67.6 | 6643 | d__Bacteria;p__Proteobacteria;c__Alphaproteobacteria;o__Rhodospirillales_B;f__Magnetospirillaceae;g__;s__ |
| nJSW_bin1 | 85.14 | 1.68 | 76.74 | 335 | 106146 | 5862517 | 42.2 | 33993 | d__Bacteria;p__Proteobacteria;c__Magnetococcia;o__Magnetococcales;f__UBA8363;g__GCA-2753565;s__ |
| nJSW_bin2 | 99.16 | 3.64 | 80.96 | 170 | 177271 | 4942843 | 55.33 | 50362 | d__Bacteria;p__Proteobacteria;c__Magnetococcia;o__Magnetococcales;f__;g__;s__ |
| nJSW_bin3 | 90.55 | 1.68 | 82.15 | 103 | 161720 | 2838557 | 44.6 | 45395 | d__Bacteria;p__Proteobacteria;c__Magnetococcia;o__Magnetococcales;f__UBA8363;g__GCA-2753565;s__ |
| nKLK_bin1 | 94.53 | 0 | 94.53 | 89 | 290812 | 3428751 | 59.34 | 60596 | d__Bacteria;p__Proteobacteria;c__Alphaproteobacteria;o__Rhodospirillales_B;f__WMHbin7;g__WMHbin7;s__ |
| nKLK_bin12 | 90.62 | 0.89 | 86.17 | 96 | 384605 | 5633167 | 32.16 | 134958 | d__Bacteria;p__Bdellovibrionota;c__Bacteriovoracia;o__Bacteriovoracales;f__Bacteriovoracaceae;g__;s__ |
| nKLK_bin4 | 80.66 | 2.94 | 65.96 | 225 | 129112 | 4028132 | 54.24 | 26216 | d__Bacteria;p__Proteobacteria;c__Magnetococcia;o__Magnetococcales;f__UBA8363;g__UBA8363;s__UBA8363 sp002753735 |
| nKLK_bin5 | 78.36 | 1.75 | 69.61 | 212 | 147700 | 4586144 | 41.45 | 36596 | d__Bacteria;p__Bdellovibrionota;c__Bacteriovoracia;o__Bacteriovoracales;f__Bacteriovoracaceae;g__;s__ |

| | | | | | | | | | |
|---|---|---|---|---|---|---|---|---|---|
| nKLK_bin6 | 90.48 | 0 | 90.48 | 88 | 377665 | 5251075 | 45.21 | 107714 | d__Bacteria;p__SAR324;c__SAR324;o__SAR324;f__;g__;s__ |
| nMBP_bin3 | 67.5 | 1.08 | 62.1 | 41 | 312199 | 1969487 | 50.38 | 106662 | d__Bacteria;p__Omnitrophota;c__koll11;o__UBA10015;f__GCA-002753745;g__GCA-2753465;s__GCA-2753465 sp002753465 |
| nMBP_bin6 | 73.26 | 1.37 | 66.41 | 481 | 65800 | 3695133 | 59.23 | 10704 | d__Bacteria;p__Proteobacteria;c__Magnetococcia;o__Magnetococcales;f__WMHbin3;g__WMHbin3;s__ |
| nMY_bin1 | 97.06 | 0.91 | 92.51 | 48 | 204131 | 2908520 | 44.36 | 104491 | d__Bacteria;p__Nitrospirota;c__Thermodesulfovibrionia;o__Thermodesulfovibrionales;f__UBA9935;g__MYbin3;s__MYbin3 sp002753335 |
| nMY_bin2 | 90.08 | 2.73 | 76.43 | 127 | 165208 | 3490049 | 49 | 45806 | d__Bacteria;p__Nitrospirota;c__Thermodesulfovibrionia;o__Thermodesulfovibrionales;f__Magnetobacteriaceae;g__Magnetobacterium;s__Magnetobacterium casensis |
| nMY_bin3 | 59.65 | 0 | 59.65 | 278 | 32451 | 1921906 | 49.89 | 8414 | d__Bacteria;p__Nitrospirota;c__Thermodesulfovibrionia;o__Thermodesulfovibrionales;f__Magnetobacteriaceae;g__Magnetobacterium;s__ |
| nMY_bin4 | 95.76 | 1.36 | 88.96 | 213 | 99834 | 3929094 | 44.04 | 34309 | d__Bacteria;p__Nitrospirota;c__Thermodesulfovibrionia;o__Thermodesulfovibrionales;f__Magnetobacteriaceae;g__Magnetobacterium;s__Magnetobacterium sp002753395 |
| nMY_bin5 | 52.67 | 0.42 | 50.57 | 376 | 31340 | 2249923 | 56.15 | 6976 | d__Bacteria;p__Proteobacteria;c__Magnetococcia;o__Magnetococcales;f__DC0425bin3;g__HA3dbin3;s__ |
| nMY_bin6 | 85.64 | 5.45 | 58.39 | 171 | 159608 | 3404042 | 47.77 | 27655 | d__Bacteria;p__Nitrospirota;c__Thermodesulfovibrionia;o__Thermodesulfovibrionales;f__Magnetobacteriaceae;g__HCH-1;s__HCH-1 sp002753305 |
| nN2-2_bin1 | 99.5 | 0.5 | 97 | 129 | 189745 | 4326850 | 65.17 | 58814 | d__Bacteria;p__Proteobacteria;c__Alphaproteobacteria;o__Rhodospirillales_B;f__Magnetospirillaceae;g__Magnetospirillum;s__Magnetospirillum moscoviense |
| nN2-2_bin2 | 76.25 | 0.5 | 73.75 | 602 | 37326 | 4106508 | 67.51 | 8405 | d__Bacteria;p__Proteobacteria;c__Alphaproteobacteria;o__Rhodospirillales_B;f__Magnetospirillaceae;g__;s__ |
| nN2-2_bin5 | 80.97 | 0.65 | 77.72 | 215 | 66927 | 3179296 | 37.93 | 18110 | d__Bacteria;p__Desulfobacterota;c__Desulfobacteria;o__Desulfobacterales;f__Desulfobacteraceae;g__Desulfamplus;s__ |
| nN3_bin14 | 70.68 | 0.75 | 66.93 | 499 | 54722 | 3735703 | 67.66 | 9763 | d__Bacteria;p__Proteobacteria;c__Alphaproteobacteria;o__Rhodospirillales_B;f__Magnetospirillaceae;g__;s__ |

| | | | | | | | | | |
|---|---|---|---|---|---|---|---|---|---|
| nN3_bin16 | 83.77 | 1.79 | 74.82 | 413 | 141348 | 4958770 | 31.53 | 18413 | d__Bacteria;p__Bdellovibrionota;c__Bacteriovoracia;o__Bacteriovoracales;f__Bacteriovoracaceae;g__;s__ |
| nN3_bin31 | 91.07 | 2.68 | 77.67 | 114 | 409610 | 5611739 | 40.9 | 99534 | d__Bacteria;p__Bdellovibrionota;c__Bacteriovoracia;o__Bacteriovoracales;f__Bacteriovoracaceae;g__;s__ |
| nNGH_bin12 | 88.82 | 1.2 | 82.82 | 358 | 52612 | 3253043 | 54.86 | 12350 | d__Bacteria;p__Nitrospinota;c__UBA7883;o__UBA7883;f__UBA7883;g__;s__ |
| nNGH_bin13 | 95.66 | 0 | 95.66 | 314 | 55101 | 3824539 | 63.09 | 18740 | d__Bacteria;p__Proteobacteria;c__Magnetococcia;o__;f__;g__;s__ |
| nNGH_bin14 | 94.12 | 2.1 | 83.62 | 358 | 97151 | 4514666 | 60.22 | 20051 | d__Bacteria;p__Proteobacteria;c__Magnetococcia;o__Magnetococcales;f__;g__;s__ |
| nNGH_bin2 | 96.47 | 3.36 | 79.67 | 133 | 144624 | 4022416 | 61.98 | 48446 | d__Bacteria;p__Proteobacteria;c__Magnetococcia;o__Magnetococcales;f__WMHbin3;g__WMHbin3;s__ |
| nPC_bin1 | 99.58 | 0.84 | 95.38 | 68 | 276500 | 2016854 | 47.46 | 56539 | d__Bacteria;p__Proteobacteria;c__Zetaproteobacteria;o__Mariprofundales;f__Mariprofundaceae;g__GCA-2753275;s__GCA-2753275 sp002753275 |
| nPCR_bin10 | 90.84 | 0 | 90.84 | 241 | 290123 | 5133600 | 42 | 50150 | d__Bacteria;p__SAR324;c__SAR324;o__SAR324;f__GCA-2753255;g__GCA-2753255;s__GCA-2753255 sp002753255 |
| nPCR_bin5 | 93.99 | 0.75 | 90.24 | 299 | 109170 | 3543883 | 34.87 | 21360 | d__Bacteria;p__Proteobacteria;c__Gammaproteobacteria;o__;f__;g__;s__ |
| nPCR_bin7 | 92.94 | 0.42 | 90.84 | 81 | 489404 | 5795165 | 44.77 | 123521 | d__Bacteria;p__SAR324;c__SAR324;o__SAR324;f__GCA-2753255;g__;s__ |
| nPCR_bin9 | 79.81 | 1.71 | 71.26 | 410 | 59515 | 2645057 | 37.05 | 7923 | d__Bacteria;p__Nitrospinota;c__UBA7883;o__UBA7883;f__;g__;s__ |
| nQXH1_bin1 | 98.51 | 0.5 | 96.01 | 23 | 869793 | 3628312 | 59.95 | 458012 | d__Bacteria;p__Proteobacteria;c__Alphaproteobacteria;o__Rhodospirillales_B;f__WMHbin7;g__WMHbin7;s__ |
| nQXH2_bin1 | 92.24 | 1.68 | 83.84 | 389 | 55544 | 3820765 | 61.35 | 19017 | d__Bacteria;p__Proteobacteria;c__Magnetococcia;o__Magnetococcales;f__WMHbin3;g__WMHbin3;s__ |

| | | | | | | | | | |
|---|---|---|---|---|---|---|---|---|---|
| nQXH2_bin5 | 98.62 | 2.52 | 86.02 | 44 | 309908 | 3842963 | 59.16 | 133818 | d__Bacteria;p__Proteobacteria;c__Magnetococcia;o__Magnetococcales;f__WMHbin3;g__WMHbin3;s__ |
| nS315_bin20 | 63.44 | 1.08 | 58.04 | 200 | 31349 | 1377317 | 56.96 | 8753 | d__Bacteria;p__Omnitrophota;c__koll11;o__2-02-FULL-51-18;f__;g__;s__ |
| nS315_bin24 | 51.04 | 0 | 51.04 | 115 | 88017 | 1495409 | 37.12 | 25313 | d__Bacteria;p__Omnitrophota;c__koll11;o__UBA1560;f__Omnitrophaceae_A;g__;s__ |
| nS315_bin3 | 84.62 | 0.16 | 83.82 | 282 | 75205 | 3601388 | 40.67 | 19749 | d__Bacteria;p__Desulfobacterota;c__Desulfobacteria;o__Desulfobacterales;f__Desulfobacteraceae;g__Desulfamplus;s__ |
| nS315_bin38 | 68.97 | 0 | 68.97 | 429 | 35700 | 2468637 | 62.18 | 6964 | d__Bacteria;p__Proteobacteria;c__Gammaproteobacteria;o__Chromatiales;f__Sedimenticolaceae;g__;s__ |
| nS315_bin44 | 88.39 | 0.32 | 86.79 | 200 | 216132 | 3860279 | 36.17 | 49168 | d__Bacteria;p__Desulfobacterota;c__Desulfobacteria;o__Desulfobacterales;f__Desulfobacteraceae;g__Desulfamplus;s__ |
| nS315_bin9 | 96.77 | 1.75 | 88.02 | 195 | 107956 | 3171500 | 37.59 | 28298 | d__Bacteria;p__Desulfobacterota;c__Desulfobacteria;o__Desulfobacterales;f__Desulfobacteraceae;g__Desulfamplus;s__ |
| nSSYD_bin4 | 94.65 | 0.33 | 93 | 213 | 99656 | 3182839 | 63.34 | 25241 | d__Bacteria;p__Proteobacteria;c__Alphaproteobacteria;o__Rhodospirillales_A;f__Magnetovibrionaceae;g__;s__ |
| nTS_bin1 | 98.71 | 1.36 | 91.91 | 210 | 176123 | 5131725 | 32.05 | 47251 | d__Bacteria;p__Desulfobacterota;c__Desulfobacteria;o__Desulfobacterales;f__YD0425bin50;g__;s__ |
| nTS_bin10 | 88.8 | 0.56 | 86 | 344 | 128004 | 5756581 | 44.87 | 28409 | d__Bacteria;p__SAR324;c__SAR324;o__SAR324;f__;g__;s__ |
| nTS_bin12 | 74.26 | 1.03 | 69.11 | 329 | 62503 | 2401499 | 63.33 | 10349 | d__Bacteria;p__Proteobacteria;c__Alphaproteobacteria;o__Rhodospirillales_A;f__Magnetovibrionaceae;g__;s__ |
| nTS_bin13 | 58.65 | 0 | 58.65 | 642 | 47405 | 3005010 | 56.47 | 5156 | d__Bacteria;p__Planctomycetota;c__SZUA-567;o__;f__;g__;s__ |
| nTS_bin15 | 73.57 | 1.94 | 63.87 | 851 | 44670 | 5493383 | 40.46 | 7452 | d__Bacteria;p__Desulfobacterota;c__Desulfobacteria;o__Desulfobacterales;f__Desulfobacteraceae;g__Desulfamplus;s__ |

| | | | | | | | | | |
|---|---|---|---|---|---|---|---|---|---|
| nTS_bin18 | 90.84 | 1.79 | 81.89 | 225 | 106673 | 3405799 | 45.02 | 24782 | d__Bacteria;p__Desulfobacterota;c__Desulfobulbia;o__Desulfobulbales;f__Desulfurivibrionaceae;g__;s__ |
| nTS_bin2 | 92.43 | 0.44 | 90.23 | 223 | 66268 | 2871709 | 52.9 | 19518 | d__Bacteria;p__Proteobacteria;c__Gammaproteobacteria;o__Thiohalomonadales;f__Thiohalomonadaceae;g__;s__ |
| nTS_bin20 | 78.06 | 5.22 | 51.96 | 402 | 69041 | 3629071 | 47.21 | 12968 | d__Bacteria;p__Desulfobacterota;c__Desulfobacteria;o__Desulfobacterales;f__Desulfobacteraceae;g__Desulfamplus;s__ |
| nTS_bin21 | 61.77 | 1.47 | 54.42 | 548 | 24652 | 2809879 | 39.13 | 6428 | d__Bacteria;p__Desulfobacterota;c__Desulfobacteria;o__Desulfobacterales;f__Desulfobacteraceae;g__Desulfamplus;s__ |
| nTS_bin4 | 97.42 | 1.29 | 90.97 | 196 | 79298 | 3232689 | 37.6 | 26935 | d__Bacteria;p__Desulfobacterota;c__Desulfobacteria;o__Desulfobacterales;f__Desulfobacteraceae;g__Desulfamplus;s__ |
| nW3_bin14 | 93.01 | 1.08 | 87.61 | 93 | 490973 | 3659045 | 49.49 | 72303 | d__Bacteria;p__Omnitrophota;c__koll11;o__UBA1560;f__Omnitrophaceae_A;g__;s__ |
| nW3_bin19 | 60.7 | 0 | 60.7 | 470 | 69206 | 4368697 | 38.41 | 14912 | d__Bacteria;p__Bdellovibrionota;c__Bacteriovoracia;o__Bacteriovoracales;f__Bacteriovoracaceae;g__;s__ |
| nW5_bin1 | 96.94 | 1.68 | 88.54 | 114 | 225186 | 4103378 | 59.91 | 70955 | d__Bacteria;p__Proteobacteria;c__Magnetococcia;o__Magnetococcales;f__WMHbin3;g__WMHbin3;s__ |
| nW5_bin3 | 88.72 | 0.84 | 84.52 | 353 | 90372 | 3790407 | 59.17 | 16624 | d__Bacteria;p__Proteobacteria;c__Magnetococcia;o__Magnetococcales;f__WMHbin3;g__WMHbin3;s__ |
| nwag_bin3 | 68.82 | 0.18 | 67.92 | 588 | 39320 | 4144413 | 41.73 | 9132 | d__Bacteria;p__Planctomycetota;c__UBA11346;o__;f__;g__;s__ |
| nwag_bin5 | 87.17 | 1.26 | 80.87 | 238 | 112061 | 3957690 | 38.49 | 22682 | d__Bacteria;p__Proteobacteria;c__Magnetococcia;o__Magnetococcales;f__UBA8363;g__GCA-2753565;s__ |
| nwal_bin5 | 78.85 | 1.59 | 70.9 | 459 | 93109 | 3818843 | 55.87 | 12313 | d__Bacteria;p__Proteobacteria;c__Magnetococcia;o__;f__;g__;s__ |
| nWMH_bin1 | 96.64 | 2.63 | 83.49 | 116 | 265640 | 4373155 | 54.97 | 83117 | d__Bacteria;p__Proteobacteria;c__Magnetococcia;o__Magnetococcales;f__WMHbin3;g__WMHbinv6;s__WMHbinv6 sp002753135 |

| | | | | | | | | | |
|---|---|---|---|---|---|---|---|---|---|
| nWMH_bin2 | 95.38 | 0.84 | 91.18 | 175 | 127019 | 3812214 | 57.17 | 39732 | d__Bacteria;p__Proteobacteria;c__Magnetococcia;o__Magnetococcales;f__DC0425bin3;g__HA3dbin3;s__ |
| nWMH_bin3 | 96.89 | 2.94 | 82.19 | 167 | 167472 | 4974826 | 61.57 | 59272 | d__Bacteria;p__Proteobacteria;c__Magnetococcia;o__Magnetococcales;f__WMHbin3;g__WMHbin3;s__WMHbin3 sp002753185 |
| nWMH_bin4 | 97.48 | 2.1 | 86.98 | 207 | 107318 | 4156077 | 54.29 | 31156 | d__Bacteria;p__Proteobacteria;c__Magnetococcia;o__Magnetococcales;f__UBA8363;g__UBA8363;s__UBA8363 sp002753735 |
| nWMH_bin5 | 97.01 | 0 | 97.01 | 80 | 521566 | 3563391 | 59.35 | 105961 | d__Bacteria;p__Proteobacteria;c__Alphaproteobacteria;o__Rhodospirillales_B;f__WMHbin7;g__WMHbin7;s__WMHbin7 sp002753155 |
| nWMH_bin6 | 73.74 | 0.91 | 69.19 | 351 | 48456 | 2327780 | 63.77 | 9028 | d__Bacteria;p__Nitrospinota;c__UBA7883;o__UBA7883;f__UBA7883;g__;s__ |
| nWRX1_bin1 | 75.18 | 1.68 | 66.78 | 200 | 58756 | 2629331 | 59.27 | 17832 | d__Bacteria;p__Proteobacteria;c__Magnetococcia;o__Magnetococcales;f__WMHbin3;g__HCHbin5;s__ |
| nWRX1_bin11 | 75.33 | 5.04 | 50.13 | 254 | 92361 | 3564446 | 60.94 | 22436 | d__Bacteria;p__Proteobacteria;c__Magnetococcia;o__Magnetococcales;f__WMHbin3;g__WMHbin3;s__ |
| nWRX1_bin12 | 58.4 | 1.26 | 52.1 | 199 | 90776 | 2774935 | 62.96 | 27522 | d__Bacteria;p__Proteobacteria;c__Magnetococcia;o__Magnetococcales;f__;g__;s__ |
| nWRX1_bin3 | 91.04 | 3.36 | 74.24 | 184 | 155931 | 3146079 | 54.88 | 35126 | d__Bacteria;p__Proteobacteria;c__Magnetococcia;o__Magnetococcales;f__;g__;s__ |
| nWRX1_bin5 | 94.12 | 2.1 | 83.62 | 173 | 132784 | 4551111 | 54.4 | 42614 | d__Bacteria;p__Proteobacteria;c__Magnetococcia;o__Magnetococcales;f__UBA8363;g__UBA8363;s__UBA8363 sp002753735 |
| nWRX1_bin6 | 83.19 | 5.04 | 57.99 | 283 | 157526 | 4081065 | 57.46 | 24368 | d__Bacteria;p__Proteobacteria;c__Magnetococcia;o__Magnetococcales;f__DC0425bin3;g__HA3dbin3;s__ |
| nWRX2_bin6 | 97.48 | 2.1 | 86.98 | 210 | 120898 | 4720416 | 54.25 | 37239 | d__Bacteria;p__Proteobacteria;c__Magnetococcia;o__Magnetococcales;f__UBA8363;g__UBA8363;s__UBA8363 sp002753735 |
| nWRX3_bin10 | 58.62 | 0 | 58.62 | 173 | 117121 | 3279066 | 62.87 | 36559 | d__Bacteria;p__Proteobacteria;c__Magnetococcia;o__Magnetococcales;f__;g__;s__ |

| | | | | | | | | | |
|---|---|---|---|---|---|---|---|---|---|
| nWRX3_bin11 | 65.43 | 0 | 65.43 | 287 | 46469 | 1964340 | 53.76 | 9873 | d__Bacteria;p__Proteobacteria;c__Magnetococcia;o__Magnetococcales;f__;g__;s__ |
| nWRX3_bin12 | 78.88 | 3.85 | 59.63 | 234 | 101072 | 3749591 | 52.93 | 22708 | d__Bacteria;p__Proteobacteria;c__Magnetococcia;o__Magnetococcales;f__UBA8363;g__UBA8363;s__ |
| nWRX3_bin2 | 95.63 | 0 | 95.63 | 197 | 276777 | 4705621 | 54.61 | 41621 | d__Bacteria;p__Proteobacteria;c__Magnetococcia;o__Magnetococcales;f__DC0425bin3;g__HA3dbin3;s__ |
| nWRX3_bin4 | 78.32 | 0.18 | 77.42 | 411 | 29825 | 2589846 | 66.09 | 8279 | d__Bacteria;p__Proteobacteria;c__Alphaproteobacteria;o__Rhodospirillales;f__Rhodospirillaceae;g__;s__ |
| nWRX3_bin6 | 97.48 | 2.94 | 82.78 | 219 | 148025 | 4250240 | 52.02 | 32202 | d__Bacteria;p__Proteobacteria;c__Magnetococcia;o__Magnetococcales;f__UBA8363;g__UBA8363;s__ |
| nWRX3_bin7 | 61.21 | 1.72 | 52.61 | 501 | 43791 | 2890474 | 61.11 | 7739 | d__Bacteria;p__Proteobacteria;c__Magnetococcia;o__Magnetococcales;f__WMHbin3;g__WMHbin3;s__ |
| nXX_bin1 | 93.7 | 0.05 | 93.45 | 228 | 125110 | 4584649 | 47.04 | 34535 | d__Bacteria;p__UBA10199;c__UBA10199;o__;f__;g__;s__ |
| nXX_bin12 | 57.14 | 0 | 57.14 | 369 | 27988 | 2585509 | 37.76 | 9633 | d__Bacteria;p__Desulfobacterota;c__Desulfobacteria;o__Desulfobacterales;f__Desulfobacteraceae;g__Desulfamplus;s__ |
| nXX_bin4 | 79.91 | 1.17 | 74.06 | 123 | 110426 | 2023619 | 46.42 | 29252 | d__Bacteria;p__Omnitrophota;c__koll11;o__UBA10015;f__GCA-002753745;g__GCA-2753745;s__ |
| nYD0423_bin2 | 96.64 | 2.1 | 86.14 | 97 | 252730 | 4218733 | 56.86 | 82296 | d__Bacteria;p__Proteobacteria;c__Magnetococcia;o__Magnetococcales;f__UBA8363;g__UBA8363;s__ |
| nYD0423_bin3 | 98.94 | 1.26 | 92.64 | 146 | 227356 | 4504451 | 55.48 | 60044 | d__Bacteria;p__Proteobacteria;c__Magnetococcia;o__Magnetococcales;f__WMHbin3;g__WMHbinv6;s__WMHbinv6 sp002753095 |
| nYD0425_bin13 | 98.1 | 1.26 | 91.8 | 231 | 135872 | 4219420 | 55.36 | 32122 | d__Bacteria;p__Proteobacteria;c__Magnetococcia;o__Magnetococcales;f__WMHbin3;g__WMHbinv6;s__WMHbinv6 sp002753095 |
| nYD0425_bin5 | 96.77 | 0.52 | 94.17 | 133 | 381417 | 5628873 | 32.38 | 103346 | d__Bacteria;p__Desulfobacterota;c__Desulfobacteria;o__Desulfobacterales;f__YD0425bin51;g__YD0425bin51;s__YD0425bin51 sp002753225 |

| | | | | | | | | | |
|---|---|---|---|---|---|---|---|---|---|
| nYD042<br>5_bin6 | 93.5 | 2.19 | 82.<br>55 | 467 | 112709 | 5417822 | 36.8<br>2 | 181<br>77 | d__Bacteria;p__Desulfobacterota;c__Desulfobacteria;o__Desulfobacterales;f__YD0425<br>bin50;g__YD0425bin50;s__YD0425bin50 sp002753105 |
| nYQH5<br>6_bin3 | 81.68 | 0.5 | 79.<br>18 | 436 | 49768 | 2796001 | 65.5<br>7 | 857<br>7 | d__Bacteria;p__Proteobacteria;c__Alphaproteobacteria;o__Rhodospirillales;f__Rhodosp<br>irillaceae;g__;s__ |

**Supplementary Table 3. Previously published MTB genomes included in this study.**

| Organism name | GenBank accession number |
|---|---|
| Alphaproteobacteria bacterium WMHbin7 | PDZW00000000 |
| Candidatus Lambdaproteobacteria bacterium PCRbin3 | PDZZ00000000 |
| Candidatus Magnetaquicoccus inordinatus strain UR-1 | RXIU00000000 |
| Candidatus Magnetobacterium bavaricum isolate TM-1 | LACI00000000 |
| Candidatus Magnetobacterium casensis strain MYR-1 | JMFO00000000 |
| Candidatus Magnetococcus massalia MO-1 | JN177318 |
| Candidatus Magnetoglobus multicellularis str. Araruama | ATBP00000000 |
| Candidatus Magnetominusculus xianensis strain HCH-1 | LNQR00000000 |
| Candidatus Magnetomorum sp. HK-1 | JPDT00000000 |
| Candidatus Magnetoovum chiemensis strain CS-04 | JZJI00000000 |
| Candidatus Omnitrophica bacterium isolate Cal1bin1 | PEAR00000000 |
| Candidatus Omnitrophica bacterium isolate MBPbin6 | PEAF00000000 |
| Candidatus Omnitrophus magneticus isolate SKK-01 | JYNY00000000 |
| Candidatus Terasakiella magnetica strain PR-1 | FLYE00000000 |
| Deltaproteobacteria bacterium isolate ER2bin7 | PEAL00000000 |
| Deltaproteobacteria bacterium YD0425bin50 | PDZT00000000 |
| Deltaproteobacteria bacterium YD0425bin51 | PDZS00000000 |
| Desulfamplus magnetovallimortis strain BW-1 | FWEV00000000 |
| Desulfovibrio magneticus RS-1 | NC_012796 |
| Ectothiorhodospiraceae bacterium BW-2 | CP032507 |
| Latescibacteria bacterium SCGC AAA252-B13 | ASWY00000000 |
| Magnetococcales bacterium DC0425bin3 | PEAP00000000 |
| Magnetococcales bacterium DCbin2 | PEAO00000000 |

| | |
|---|---|
| Magnetococcales bacterium DCbin4 | PEAN00000000 |
| Magnetococcales bacterium HA3dbin3 | PEAJ00000000 |
| Magnetococcales bacterium HCHbin5 | PEAG00000000 |
| Magnetococcales bacterium isolate ER1bin7 | PEAM00000000 |
| Magnetococcales bacterium isolate HA3dbin1 | PEAK00000000 |
| Magnetococcales bacterium isolate HAa3bin1 | PEAI00000000 |
| Magnetococcales bacterium isolate WMHbin1 | PDZY00000000 |
| Magnetococcales bacterium isolate YD0425bin7 | PDZU00000000 |
| Magnetococcales bacterium WMHbin3 | PDZX00000000 |
| Magnetococcales bacterium WMHbinv6 | PDZV00000000 |
| Magnetococcus marinus MC-1 | NC_008576 |
| Magnetofaba australis IT-1 | LVJN00000000 |
| Magnetospira sp. QH-2 | FO538765 |
| Magnetospirillum caucaseum strain SO-1 | AONQ00000000 |
| Magnetospirillum gryphiswaldense strain MSR-1 | NC_023065 |
| Magnetospirillum kuznetsovii strain LBB-42 | PGTO01000000 |
| Magnetospirillum magneticum strain AMB-1 | NC_007626 |
| Magnetospirillum magnetotacticum MS-1 | JXSL00000000 |
| Magnetospirillum marisnigri strain SP-1 | LWQT00000000 |
| Magnetospirillum moscoviense strain BB-1 | LWQU00000000 |
| Magnetospirillum sp. ME-1 | CP015848 |
| Magnetospirillum sp. XM-1 | LN997848 |
| Magnetovibrio blakemorei strain MV-1 | MCGG00000000 |
| Nitrospira bacterium SG8_35_4 | LJTM00000000 |
| Nitrospirae bacterium isolate DC0425bin1 | PEAQ00000000 |

| | |
|---|---|
| Nitrospirae bacterium isolate HCHbin1 | PEAH00000000 |
| Nitrospirae bacterium isolate MYbin2 | PEAE00000000 |
| Nitrospirae bacterium MYbin3 | PEAD00000000 |
| Nitrospirae bacterium MYbin6 | PEAC00000000 |
| Nitrospirae bacterium MYbinv3 | PEAB00000000 |
| Omnitrophica WOR_2 bacterium GWA2_45_18 | MHFX01000000 |
| Omnitrophica WOR_2 bacterium GWC2_45_7 | MHGD01000000 |
| Planctomycetes bacterium SM23_25 | LJTY00000000 |
| Terasakiella sp. SH-1 | CP038255 |
| Uncultured Desulfobacteraceae bacterium isolate CR-1 | CAACVI000000000 |
| Zetaproteobacteria bacterium isolate PCbin4 | PEAA00000000 |
