## Supplementary figures and images for "Expanding magnetic organelle biogenesis in the domain *Bacteria*"

### Supplementary Figure 1

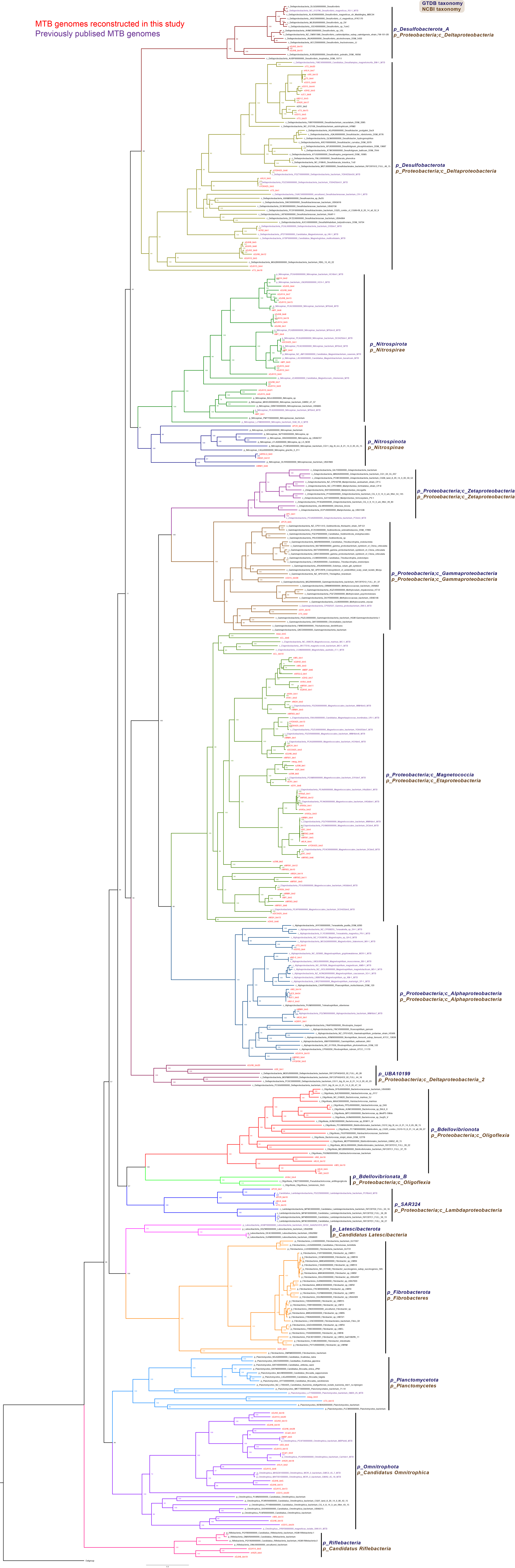
